# Accessible and reproducible mesoscale fMRI at 5.0 T: A Pulseq-based open framework for human laminar mapping

**DOI:** 10.64898/2026.02.03.703466

**Authors:** Yanting Zhu, Mengqi Jiang, Jianglian Chen, Fangyi Hao, Xin Li, Yiyun Qi, Yue Zhang, Haohao Peng, Yuhai Xie, Jiayu Zhu, Zhiwei Ma

**Author notes:** These authors contributed equally to this work (Yanting Zhu, Mengqi Jiang, Jianglian Chen, Fangyi Hao). Corresponding author: Zhiwei Ma.

## Abstract

Mesoscale, layer-specific functional MRI (fMRI) enables noninvasive access to cortical microcircuitry, yet widespread adoption has been constrained by a reliance on ultra-high field (≥7.0 T) systems and proprietary pulse sequences. To bridge this gap and enhance accessibility, we developed an open-source framework at 5.0 T for mapping laminar brain activity. This framework integrates a Pulseq-based 3D vascular space occupancy (VASO) sequence with an end-to-end data acquisition and analysis pipeline. At matched sub-millimeter resolution (0.8 mm in-plane), the Pulseq-based 3D implementation increased slab coverage by ∼1.82-fold and improved temporal signal-to-noise ratio by ∼1.50-fold relative to a vendor-provided 2D-VASO sequence. Validated using a finger-tapping paradigm, individual cerebral blood volume-weighted (VASO) laminar activation profiles consistently revealed the canonical “double-peak” pattern, with distinct superficial and deep peaks in the primary motor cortex. These profiles exhibited excellent cross-visit reliability (r = 0.80), and peak depths showed good spatial reliability (ICC = 0.69 for deep layers; ICC = 0.58 for superficial layers). Between-subject reproducibility was high (r = 0.86). Deploying the identical Pulseq protocol at an independent imaging site reproduced the characteristic double-peak laminar profiles (r = 0.63). At the group level, 5.0 T laminar profiles closely matched established 7.0 T findings, robustly resolving both deep and superficial peaks despite the lower field strength. Notably, for each participant, a single 13-minute VASO run was sufficient to resolve reliable laminar activation patterns that exhibited high consistency with multi-run averages (r = 0.78), highlighting the potential for high-throughput population studies or clinical research settings. The Pulseq-based 3D VASO sequence file, image reconstruction pipeline, and data analysis scripts are openly available to facilitate the adoption of this framework. This work establishes a practical route towards more accessible and reproducible mesoscale fMRI for studying human laminar functional architecture.

## Introduction

Cortical computation emerges from laminar input-output motifs. In the primary motor cortex (M1), superficial layers integrate thalamocortical and corticocortical inputs, whereas deep layers generate outputs to both the thalamus (corticothalamic) and downstream motor targets (corticospinal) [1]. Mapping these layer-specific processes noninvasively in humans is central to bridging systems and circuit neuroscience, enabling the investigation of canonical microcircuit models such as predictive coding frameworks and hierarchical processing theories [2]. While mesoscale neuroimaging, particularly layer-specific functional MRI (layer-fMRI), can provide sub-millimeter, depth-resolved access to cortical computations, widespread adoption remains constrained by methodological and translational barriers. These include vascular biases in blood-oxygenation-level-dependent (BOLD) contrast that conflate neural activity with venous contamination [1, 3–8], limited portability of advanced pulse sequences across vendors and sites [9, 10], and restricted availability of ultra-high-field scanners that confine deployment to specialized imaging centers [11, 12]. To broaden community access while preserving the laminar sensitivity essential for mechanistic insights, a systematic translation of layer-fMRI methods to more accessible field strengths and open-source, vendor-agnostic sequence implementations is required.

Current laminar neuroimaging methods face a trade-off between sensitivity and accessibility. Ultra-high-field (≥7.0 T) systems offer superior laminar discrimination, but high infrastructure and operating costs, together with the relatively small number of installed systems, have restricted widespread deployment [11, 12]. Conversely, recent work has demonstrated cerebral blood volume (CBV)-weighted layer-fMRI at 3.0 T [13, 14], but these approaches remain sensitivity-limited at sub-millimeter resolution and therefore often require additional averaging and denoising to achieve robust laminar profiles [13, 15, 16]. The emerging 5.0 T platform provides a compelling intermediate field strength: it delivers higher intrinsic signal-to-noise ratio (SNR) than 3.0 T, while remaining more viable for broader deployment than ultra-high-field systems [17], offering a pragmatic route toward wider adoption of mesoscale fMRI.

Despite the promise of 5.0 T, the field lacks an end-to-end, open-source 5.0 T layer-fMRI framework validated for community adoption. While 3D vascular space occupancy (VASO) sequences are well-established at ultra-high-field for layer-specific CBV mapping [1, 7, 8, 18–21], systematic implementation and validation at 5.0 T using an open-source, vendor-agnostic sequence definition remain unreported to our knowledge. A recent 5.0 T demonstration has relied on 2D-VASO, whose limited coverage restricts laminar analyses [22]. Furthermore, this work has utilized specialized radiofrequency (RF) hardware, which can complicate reproducibility and limit multi-site deployment for sites using only standard vendor-supplied coils [22]. Collectively, these limitations hinder the transition from proof-of-concept studies to a reproducible and accessible layer-fMRI framework deployable in large-scale neuroimaging and clinical studies.

We addressed these limitations by developing an open, Pulseq-based layer-fMRI framework at 5.0 T, integrating a vendor-agnostic 3D-VASO acquisition with a dedicated reconstruction and analysis pipeline (Fig 1). We benchmarked performance against a vendor-provided 2D-VASO sequence and validated laminar profiles in M1 using a finger-tapping paradigm. We further evaluated the robustness of M1 laminar profiles across longitudinal visits, between participants, and across imaging sites, and tested whether reliable canonical laminar profiles can be obtained from single 13-minute acquisitions to support high-throughput study designs. All sequence definitions and processing scripts are openly available to facilitate broader adoption of 5.0 T layer-fMRI in neuroscience and clinical research settings.

**Fig 1.**
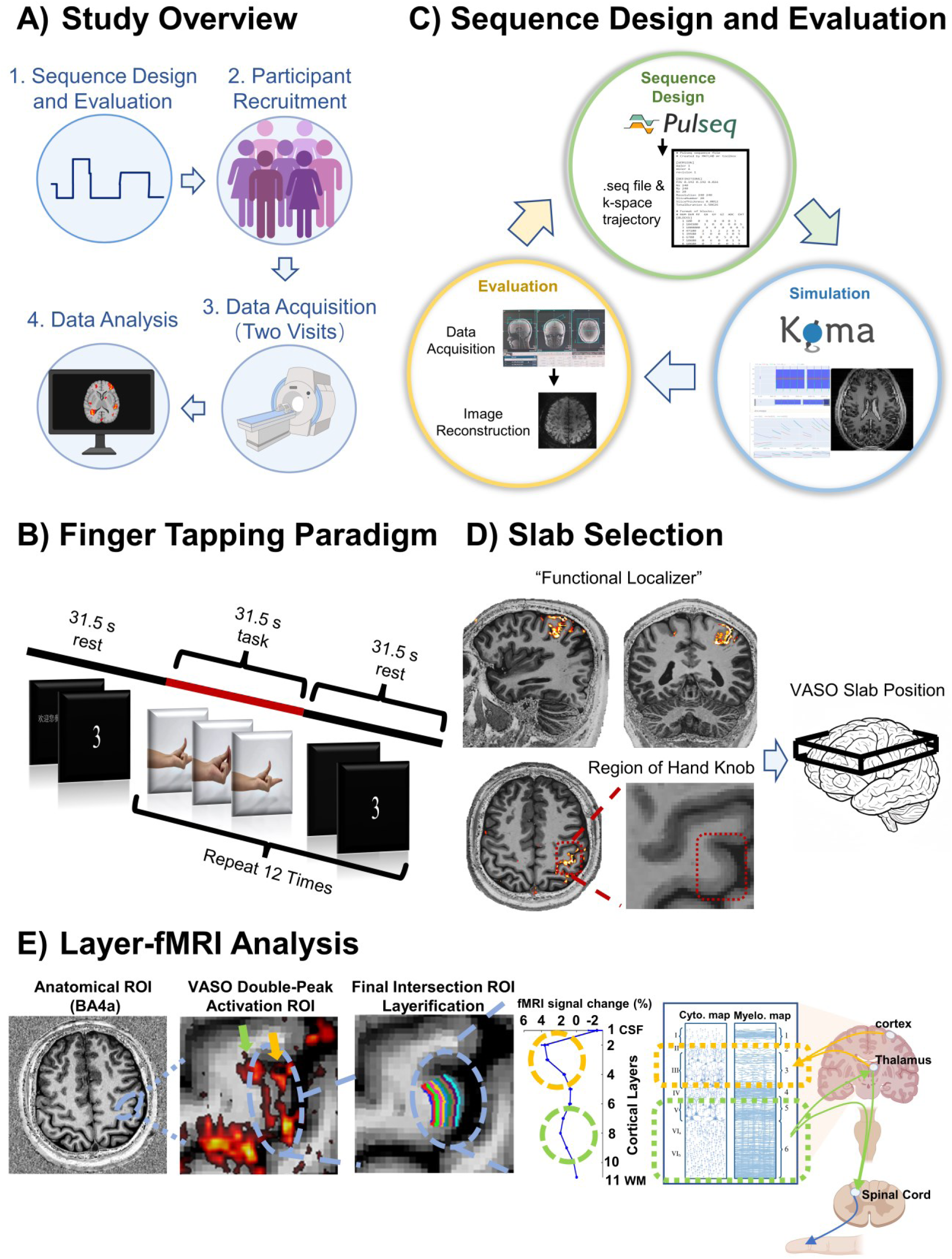
Overview of the Pulseq-based 5.0 T layer-fMRI framework. **(A)** Study workflow illustrating the pipeline from Pulseq-based sequence design to participant recruitment, multi-visit fMRI data acquisition, and layer-fMRI analysis. **(B)** Task paradigm for VASO fMRI data acquisition. Each VASO run comprised a 31.5 s pre-stimulus baseline followed by 12 alternating task and rest blocks (31.5 s each). **(C)** End-to-end workflow of Pulseq-based sequence implementation: sequence design generates vendor-agnostic .seq files; KomaMRI simulation validates sequence logic via Bloch equation modeling; MRI data acquisition collects raw data; and offline image reconstruction applies Nyquist ghost correction and GRAPPA to yield final images. **(D)** VASO slab positioning. Left: Functional localization of the M1 hand-knob using a finger-tapping task across three orthogonal planes, with a magnified view of the hand-knob anatomical landmark. Right: 3D slab placement over the motor cortex, guided by anatomical landmarks and functional activation. **(E)** Layer-fMRI analysis pipeline. ROI definition was based on VASO activation within BA4a. Cortical depth segmentation (LayNii) yielded 11 equidistant bins (L1–L11, CSF to WM). The resulting profile reflects M1 mesoscale circuitry: superficial layers (yellow) receive cortical and thalamic inputs; deep layers (green) provide output to the thalamus and spinal cord. **Abbreviations:** GRAPPA, generalized autocalibrating partially parallel acquisitions; VASO, vascular space occupancy; BOLD, blood-oxygenation-level-dependent; ROI, region of interest; BA4a, Brodmann area 4a; CSF, cerebrospinal fluid; WM, white matter.

## Results

### Pulseq-based 3D-VASO at 5.0 T substantially expands coverage (∼1.82×) with higher tSNR (∼1.50×) relative to vendor 2D-VASO

We implemented a customized Pulseq-based 3D-VASO sequence at 5.0 T (Fig 2A). Our 5.0 T 3D-VASO implementation achieved spatial and temporal acquisition parameters comparable to standard layer-fMRI protocols at 7.0 T. Fig 2B shows cortical boundaries from the MP2RAGE anatomical reference overlaid on the reconstructed VASO images from this 3D sequence, demonstrating good anatomical alignment at sub-millimeter resolution. To evaluate performance gains for fMRI, we compared this open-source Pulseq-based 3D-VASO sequence with a vendor-provided 2D-VASO sequence at a matched sub-millimeter resolution (0.8 × 0.8 × 1.2 mm^3^) within the same visit in the same participant. Quantitative analysis revealed that our 3D sequence achieved an approximately 1.82-fold increase in slab coverage and a 1.50-fold enhancement in temporal signal-to-noise ratio (tSNR) relative to the 2D sequence (Mann-Whitney U test, p < 0.0001; Fig 2C). Critically, average signal time courses extracted from M1 activation regions confirmed that the 3D-VASO signal exhibited the characteristic negative change inversely mirroring the positive BOLD response, with both signals accurately tracking the temporal dynamics of the task block design (Fig 2D). The combination of improved spatial coverage, higher tSNR, and validated CBV responses demonstrates that this open-source 3D VASO sequence constitutes a reliable tool for sub-millimeter fMRI studies at 5.0 T.

**Fig 2.**
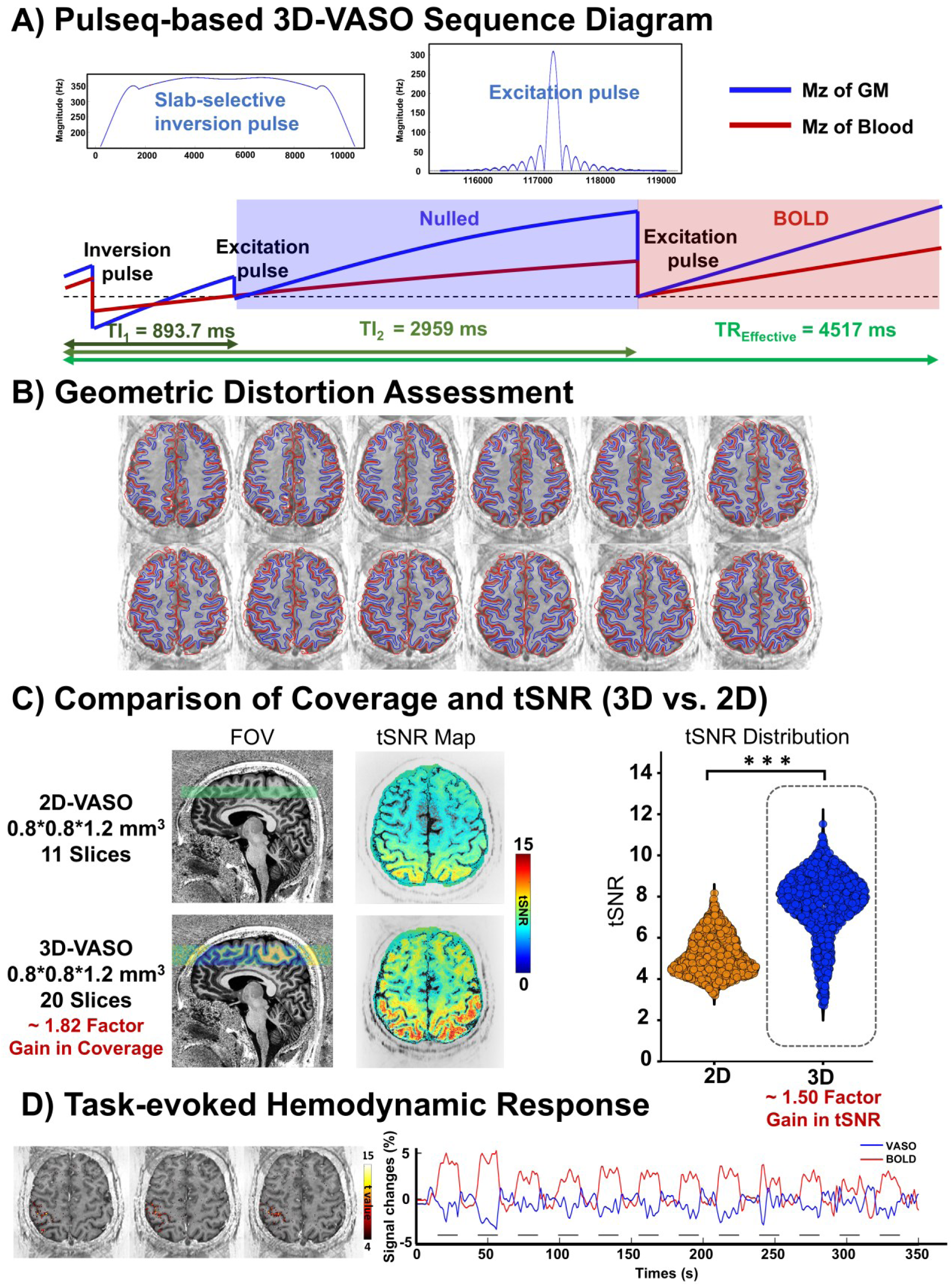
Pulseq-based 3D-VASO sequence design and benchmarking against vendor 2D-VASO. **(A)** Pulseq-based 3D-VASO sequence diagram. A TR-FOCI pulse precedes each image acquisition module. Blood-nulled (VASO) and BOLD-weighted volumes are acquired alternately. Inset: Detail of the slab-selective inversion and excitation pulses. **(B)** Geometric distortion assessment. Twelve representative slices showing cortical boundaries from the MP2RAGE anatomical image overlaid on VASO images. Red: CSF–GM boundary; blue: GM–WM boundary. **(C)** Comparison of coverage and tSNR (3D vs. 2D). Left: Sagittal images illustrating slab coverage (green overlay; ∼1.82-fold increase for 3D at matched TR; voxel size: 0.8 × 0.8 × 1.2 mm³). Middle: tSNR spatial maps. Right: Violin plots of voxelwise tSNR in the hand-knob ROI (orange: 2D; blue: 3D). The 3D sequence yielded significantly higher tSNR (p < 0.0001, Mann–Whitney U test). **(D)** Task-evoked hemodynamic responses. Percent signal change time courses averaged over motor-active voxels (t > 4; sign-flipped). VASO (blue curve) shows negative signal change reflecting increased CBV; BOLD (red curve) shows positive signal change. Gray lines indicate task block durations. **Abbreviations:** VASO, vascular space occupancy; TR-FOCI, time-resampled frequency-offset corrected inversion; BOLD, blood-oxygenation-level-dependent; MP2RAGE, magnetization prepared 2 rapid acquisition gradient echoes; CSF, cerebrospinal fluid; GM, gray matter; WM, white matter; tSNR, temporal signal-to-noise ratio; CBV, cerebral blood volume.

### Laminar VASO profiles at 5.0 T exhibit a canonical double-peak pattern in M1 with robust test–retest reliability, between-participant generalizability, and cross-site portability

Using a finger-tapping paradigm, layer-specific activation profiles were measured in M1 across longitudinal visits, participants, and two imaging sites to assess the reproducibility of laminar functional signals. In total, we analyzed 11 participants: 10 participants at Site 1 completed two imaging visits, and one participant at Site 2 completed one imaging visit. In all sessions, finger tapping evoked consistent task-related activation patterns. VASO maps showed spatially focal CBV-weighted responses in the M1 hand-knob region, whereas BOLD activation extended broadly into the surrounding sulci (Fig 3A).

**Fig 3.**
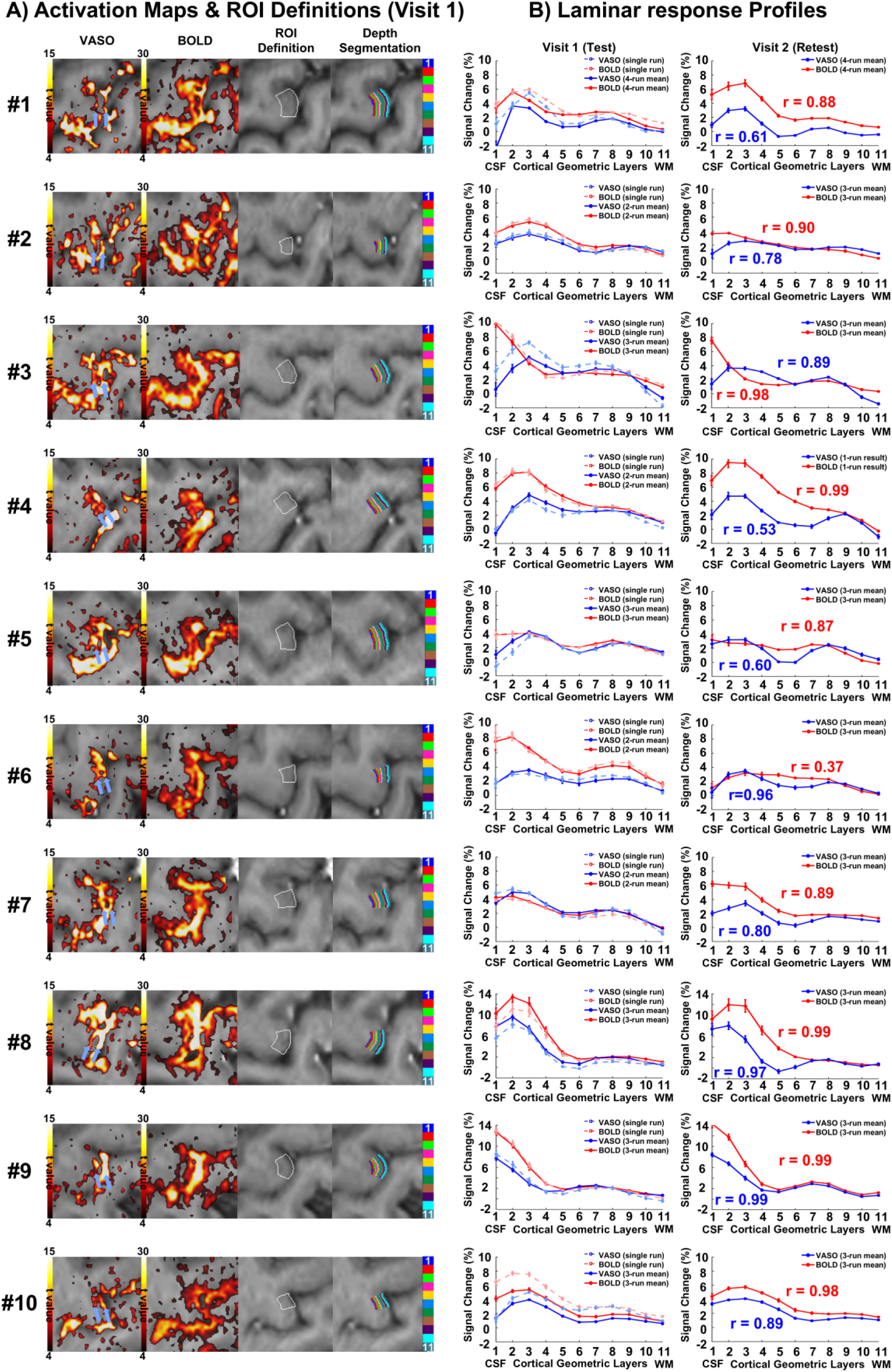
Individual-level VASO and BOLD activation maps and test-retest laminar profiles. **(A)** Activation maps and ROI definitions (Visit 1). Columns display individual-level VASO (t > 4, sign-flipped; column 1) and BOLD (t > 4; column 2) activation maps, the ROI of the lateral hand-knob area (BA4a; white outline in column 3), and cortical depth segmentation (11 equidistant bins, L1–L11, from CSF to WM; column 4). Blue arrows in VASO activation maps indicate the double-peak pattern (two depth-separated activation bands), whereas BOLD activation extends broadly into surrounding sulci. **(B)** Laminar response profiles. Plots are organized by visit: Visit 1 (Test) column compares single-run profiles (dashed lines) with multi-run averaged profiles (solid lines; e.g., “4-run mean”). Visit 2 (Retest) column displays the multi-run averaged profiles for the second visit, with Pearson correlation coefficients (r) relative to Visit 1. Curves represent mean ± SE across voxels within the ROI. Blue: VASO; red: BOLD. VASO profiles are sign-inverted for visualization. Data from all ten Site 1 participants are shown row by row. **Abbreviations:** VASO, vascular space occupancy; BOLD, blood-oxygenation-level-dependent; ROI, region of interest; BA4a, Brodmann area 4a; CSF, cerebrospinal fluid; WM, white matter; SE, standard error.

We extracted layer-specific profiles across 11 equidistant cortical depth bins, revealing distinct laminar patterns for VASO and BOLD. Based on the canonical laminar architecture of M1, in which superficial layers receive inputs from the cortex and thalamus and deep layers provide output to the thalamus and spinal cord, two distinct activity peaks at different cortical depths are expected during motor tasks [1] (Fig 1E). Consistent with this model, VASO profiles exhibited a characteristic double-peak pattern: a larger peak of 4.81% ± 2.00% signal change in superficial layers (depth bin 2.50 ± 0.69), and a smaller peak of 2.10% ± 0.66% in deep layers (depth bin 8.10 ± 0.55). The two peaks were separated by approximately 5.60 ± 0.60 depth bins, and the deep-to-superficial peak amplitude ratio for VASO was 0.48 ± 0.18 (Fig 3B). In contrast, BOLD profiles were dominated by large superficial signals of 7.47% ± 3.48% (depth bin 2.05 ± 0.83).

To quantify test–retest reliability across visits (Site 1; n = 10 participants with two visits), we compared within-participant laminar depth profiles between visits. VASO and BOLD activation maps each exhibited high spatial consistency across visits (Figs 3 and S1). VASO depth profiles demonstrated high consistency, with a mean cross-visit correlation of r = 0.80 ± 0.17 (median = 0.85), while BOLD profiles yielded r = 0.89 ± 0.19 (median = 0.94). Notably, both VASO and BOLD profiles demonstrated strong consistency (r > 0.6) in 90% of participants (Fig 3B). Peak position reproducibility was moderate (ICC = 0.58, κ_w_ = 0.56) for the VASO peak in superficial layers, and good (ICC = 0.69, κ_w_ = 0.67) for the peak in deep layers (Fig 4A).

**Fig 4.**
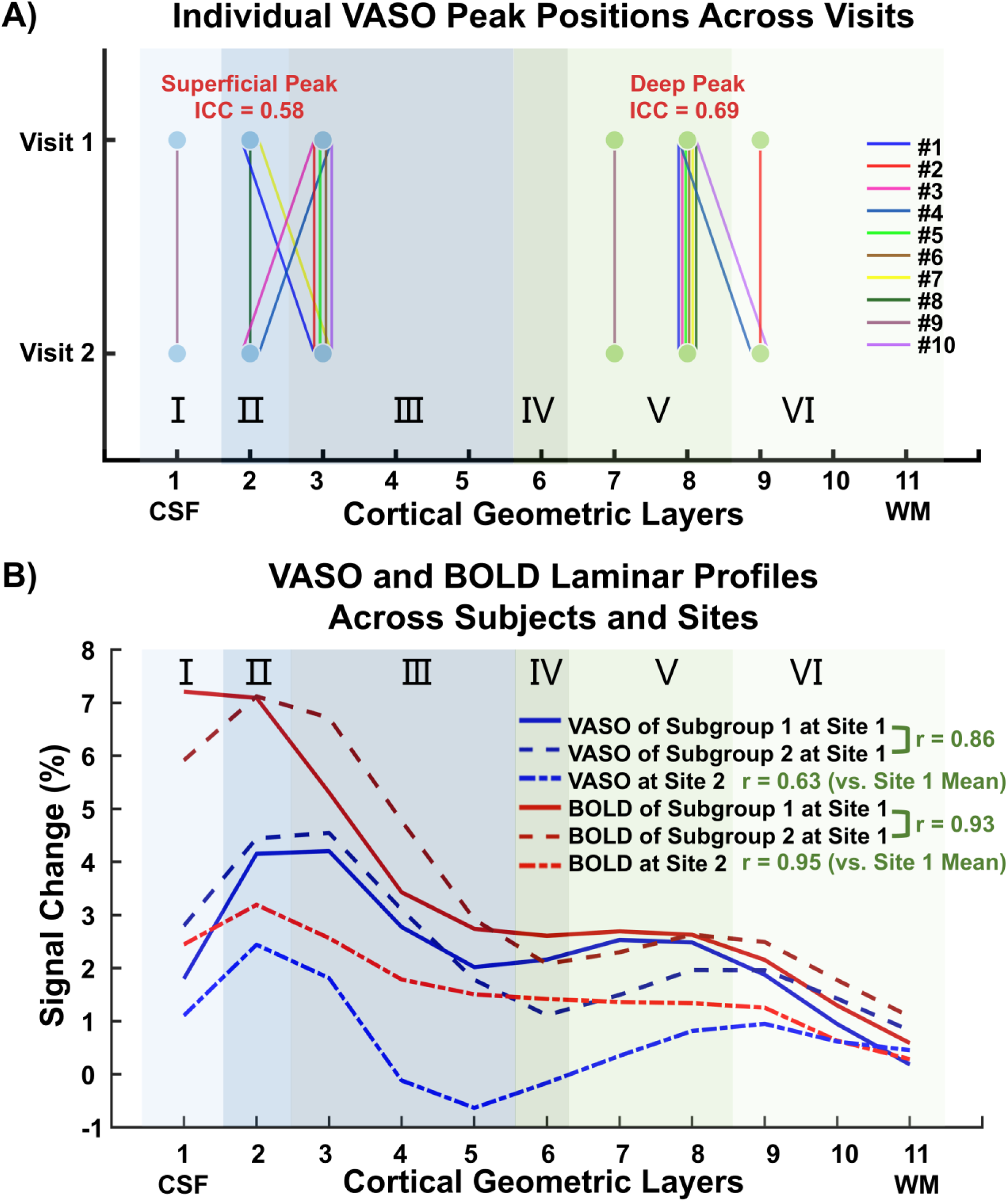
Reliability of peak positions and generalizability of laminar profiles. **(A)** Individual VASO peak positions across visits. Cortical depths of VASO signal peaks in superficial and deep layers are shown for each Site 1 participant (color-coded), with lines connecting Visit 1 and Visit 2 to illustrate test-retest correspondence. **(B)** Generalizability and portability of laminar profiles. Site 1 participants were randomly divided into two subgroups (n = 5 each). Shown are group-averaged profiles for the two subgroups and the Site 2 participant (n = 1). Upper right: Pearson correlation coefficients (r) are displayed alongside the legend: the bracketed r values indicate the consistency between the two Site 1 subgroups (generalizability), while the r values for Site 2 indicate the correlation between the Site 2 participant and the Site 1 group mean (portability). Blue curve: VASO; red curve: BOLD. VASO profiles are sign-inverted for visualization. Background shading indicates putative cytoarchitectonic layers derived from the BigBrain 3D histological atlas [69], projected onto the 11 equidistant geometric layers to visualize laminar correspondence. **Abbreviations:** VASO, vascular space occupancy; BOLD, blood-oxygenation-level-dependent.

To assess between-participant generalizability (Site 1; n = 10), participants were randomly divided into two subgroups and subgroup-averaged laminar profiles were compared. Subgroup-averaged profiles remained highly consistent (VASO: r = 0.86; BOLD: r = 0.93; Fig 4B), demonstrating that laminar activation patterns are reproducible across subgroups.

Finally, to assess cross-site portability, the identical sequence protocol was deployed at an independent site (Site 2; identical 5.0 T MRI; n = 1 participant, single visit). In this cross-site demonstration, the VASO spatial activation pattern matched Site 1 (Fig S2), and the characteristic double-peak profile was reproduced, with between-site profile correlations of 0.63 for VASO and 0.95 for BOLD (Fig 4B).

Together, these findings demonstrate that the 5.0 T Pulseq-based framework robustly captures the characteristic M1 double-peak pattern with excellent test–retest reliability (r = 0.80), strong between-participant generalizability (r = 0.86), and successful cross-site portability (r = 0.63).

### Group-averaged laminar profiles at 5.0 T replicate canonical 7.0 T double-peak patterns

To assess whether 5.0 T laminar profiles match established findings at ultra-high field, layer-specific profiles were averaged across all participants and compared quantitatively with published 7.0 T results [1]. For each participant, layer profiles were averaged across multiple runs prior to group averaging. The resulting group-mean VASO profile exhibited the characteristic double-peak pattern, showing a larger peak of 4.21% ± 0.44% signal change in superficial layers (depth bin 2; putative cytoarchitectonic layer II) and a smaller peak of 1.98% ± 0.15% in deep layers (depth bin 8; putative cytoarchitectonic layer V) (Fig 5A). Statistical analysis using a linear mixed-effects model confirmed the robustness of this double-peak feature in the VASO profile: Both the superficial peak (bins 1–3) and the deep peak (bins 7–9) exhibited significantly higher activation amplitudes compared to the intermediate and boundary layers (bins 4–6, 10–11; Superficial vs. Reference: p < 0.001; Deep vs. Reference: p < 0.05) (Fig 5B). In contrast, BOLD profiles were characterized by a single dominant superficial peak, where only the superficial peak exceeded the others (p < 0.001), while the deep peak did not differ significantly from the others (p > 0.05) (Fig 5B), reflecting BOLD’s greater sensitivity to large draining veins in superficial layers.

**Fig 5.**
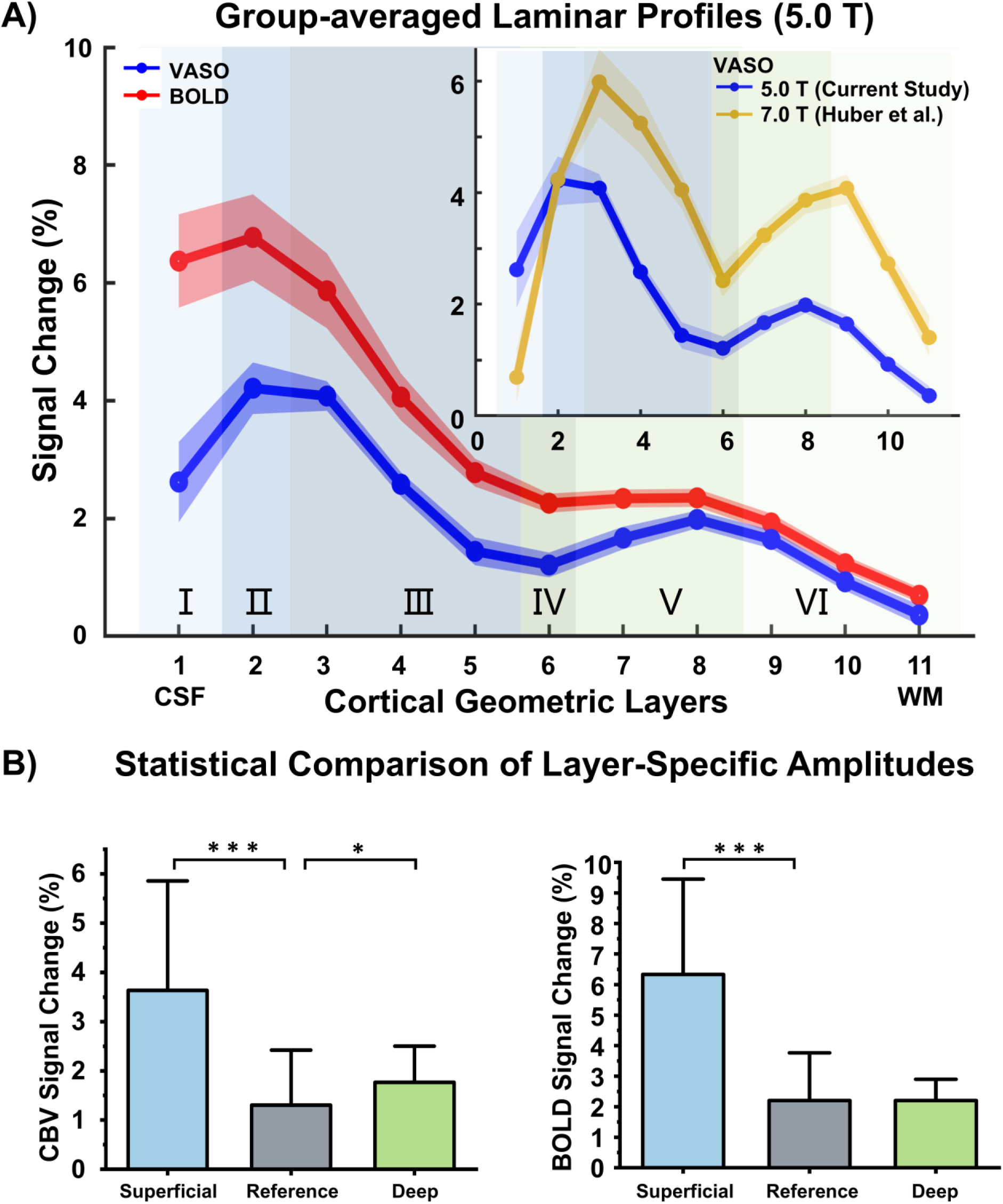
Group-level laminar profiles and statistical validation of the double-peak pattern. **(A)** Group-averaged VASO and BOLD laminar profiles (Site 1; N = 10 participants, 56 runs total). Solid lines: mean percent signal change; shaded ribbons: SE across participants. VASO profiles are sign-inverted for visualization. Inset: Comparison of 5.0 T VASO results with published 7.0 T VASO data from Huber et al. [1] (resampled to 11 depth bins). Background shading indicates putative cytoarchitectonic layers as in Fig 4. **(B)** Statistical validation of the VASO double-peak pattern. Signal amplitudes grouped by cortical depth: Superficial (bins 1–3), Deep (bins 7–9), and Reference (bins 4–6, 10–11). Bars represent mean percent signal change; error bars indicate SE. Statistical differences relative to the Reference group were assessed using a linear mixed-effects model. *p < 0.05, ***p < 0.001. **Abbreviations:** VASO, vascular space occupancy; BOLD, blood-oxygenation-level-dependent; SE, standard error.

Quantitative comparisons with published 7.0 T benchmarks confirmed that, despite an expected reduction of approximately 2 percentage points in absolute signal amplitude, the canonical double-peak pattern was preserved at 5.0 T (5.0 T VASO superficial layer: 4.21% ± 0.44%, deep layer: 1.98% ± 0.15%; 7.0 T VASO superficial layer: 6.11% ± 0.65%, deep layer: 4.12% ± 0.27%) [1] (Fig 5A, inset). These group-averaged profiles demonstrate that the 5.0 T framework robustly captures the canonical 7.0 T double-peak pattern, validating the capacity of this framework to resolve mesoscale functional architecture previously reserved for ultra-high field systems.

### Single 13-minute VASO acquisition yields reliable laminar activation profiles

To assess the feasibility of this framework for high-throughput studies and clinical settings, single functional runs were analyzed to determine whether layer-specific double-peak profiles could be robustly detected with only a single 13-minute run comprising 12 task blocks. Qualitatively, single-run VASO activation maps exhibited double-peak patterns comparable to those observed in the multi-run averages (Fig S3). Quantitatively, single-run profiles closely matched the high-tSNR reference profiles derived from the average of the remaining runs, yielding strong correlations across all participants: r = 0.78 ± 0.13 for VASO and r = 0.96 ± 0.04 for BOLD. Regarding peak spatial reliability, the superficial VASO peak demonstrated excellent consistency (ICC = 0.89, κ_w_ = 0.88), while the peak in deep layers showed moderate consistency (ICC = 0.59, κ_w_ = 0.56) (Fig 3B).

The group-averaged single-run VASO profile preserved the canonical double-peak pattern. The peak in superficial layers (depth bin 3; putative cytoarchitectonic layer III) exhibited a mean signal change of 4.76% ± 0.48%, while the peak in deep layers (depth bin 8; putative cytoarchitectonic layer V) showed 2.37% ± 0.28% (Fig 6). This group-averaged single-run profile was highly consistent with the group-averaged high-tSNR reference profile (derived from averaging the remaining runs across participants; r = 0.99), confirming that the double-peak pattern can be robustly detected from single-run acquisitions. Similarly, the group-averaged single-run BOLD profile (r > 0.99) also reliably captured the characteristic superficial-dominant activation pattern, demonstrating high signal quality across both contrasts.

**Fig 6.**
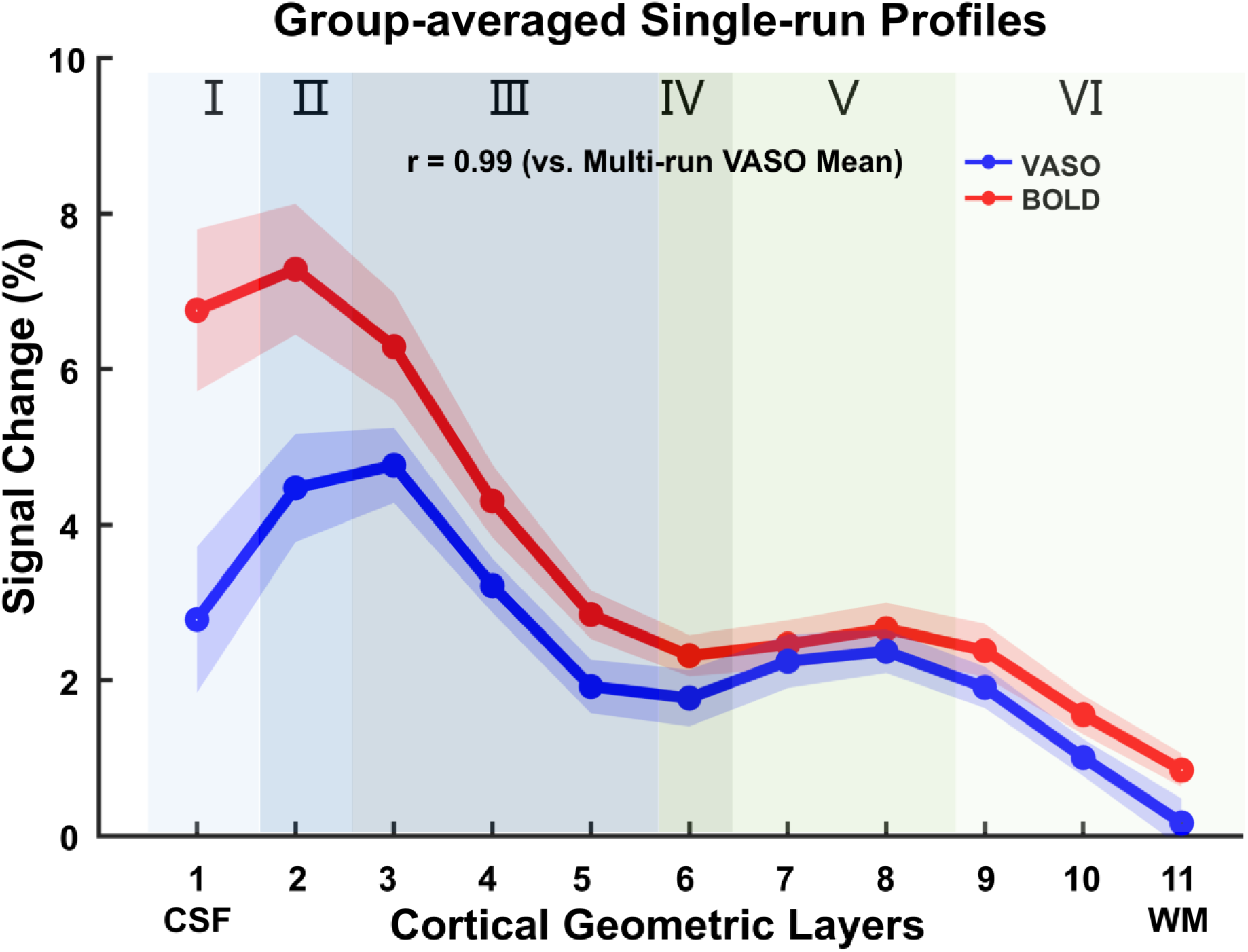
Robustness of single-run laminar profiles. Group-averaged response profiles calculated using only the first 13-minute functional run of Visit 1 (n = 10 participants). Solid lines: mean single-run response for VASO (blue curve) and BOLD (red curve); shaded areas: SE. VASO profiles are sign-inverted for visualization. Background shading indicates putative cytoarchitectonic layers as in Fig 4. The single-run VASO profile correlates strongly with the multi-run reference (r = 0.99), confirming that the canonical double-peak pattern is preserved in single-run data. **Abbreviations:** VASO, vascular space occupancy; BOLD, blood-oxygenation-level-dependent; SE, standard error.

## Discussion

In this study, we validated an open-source, Pulseq-based 5.0 T layer-fMRI framework that integrates a vendor-agnostic 3D-VASO acquisition with an end-to-end reconstruction and analysis pipeline. Benchmarking against a vendor-provided 2D sequence demonstrated substantial improvements in spatial coverage and signal quality at sub-millimeter resolution. Using a finger-tapping paradigm in M1, the framework robustly detected the canonical double-peak laminar profiles across visits, participants, and imaging sites. Quantitative comparison confirmed that 5.0 T laminar profiles closely matched published 7.0 T benchmarks. Critically, reliable layer-specific signatures can be obtained from single 13-minute acquisitions, establishing practical feasibility for high-throughput and clinical applications. By openly releasing the complete framework including pulse sequence, reconstruction and analysis scripts, we provide a standardized solution to facilitate broader adoption of layer-fMRI at this more accessible field strength.

### 5.0 T bridges the gap between sensitivity and accessibility for layer-fMRI

Achieving the sub-millimeter resolution required for layer-fMRI presents a unique challenge: it demands substantially higher intrinsic SNR than standard neuroimaging, yet widespread adoption is restricted by the limited availability and high infrastructure burden of ultra-high-field systems. 5.0 T MRI addresses this dilemma by providing substantially higher intrinsic SNR than 3.0 T, while mitigating some of the RF (B1+) inhomogeneity challenges common at 7.0 T [17, 23]. These technical characteristics place 5.0 T in a practical intermediate regime for mesoscale mapping. Both 3.0 T and 5.0 T layer-fMRI benefit substantially from advanced denoising techniques such as NORDIC [13–15]. However, the lower intrinsic SNR at 3.0 T leaves less margin for experimental imperfections—small deviations in inversion time optimization, slice positioning, or head motion can more readily compromise data quality. The higher baseline SNR at 5.0 T provides additional robustness, making single-participant laminar profiling more reliably achievable even under slightly suboptimal experimental conditions. Crucially, despite a reduction in absolute response amplitude expected with lower field strength, our group-level results demonstrate that 5.0 T robustly preserves the canonical double-peak structure reported at 7.0 T, reinforcing 5.0 T as a practical intermediate platform suitable for robust laminar mapping.

Beyond sensitivity, 5.0 T offers operational advantages that can improve the accessibility of layer-fMRI. Although some 7.0 T platforms have obtained clinical clearances, broader deployment remains limited by installation cost, siting requirements, and operational complexity. By contrast, 5.0 T systems have received regulatory clearance for whole-body clinical imaging in some jurisdictions, supporting broader translational use cases. The rapidly expanding accessibility of the 5.0 T platform, evidenced by a total installed base of over 40 systems in China alone as of 2025, underscores its potential for large-scale deployment. Furthermore, patient tolerance studies indicate high acceptance rates (> 96%) with minimal discomfort [24], largely addressing the acoustic noise and specific absorption rate (SAR) concerns often associated with ultra-high-field systems. Together, these factors support 5.0 T as a feasible platform for expanding access to high-resolution laminar imaging beyond a small number of ultra-high-field centers.

### Open-source Pulseq framework promotes standardization, portability, and accessibility of layer-fMRI

Pulseq provides an open, text-based, vendor-agnostic framework for MRI pulse-sequence development, enabling standardized sequence definitions that can be shared across platforms. It has been applied to implement mesoscale functional laminar imaging, such as 3D stack-of-spirals VASO, to improve sensitivity and specificity in high-resolution fMRI experiments [25], as well as other advanced contrasts including diffusion-weighted imaging [26] and chemical exchange saturation transfer MRI [27]. This breadth highlights Pulseq’s versatility for functional and quantitative MRI and its ability to reduce dependence on vendor-specific programming.

Pulseq enhances standardization in multi-vendor and multi-site studies. In our work, the standardized Pulseq sequence definition enabled deployment at an independent imaging site, achieving the cross-site portability of laminar profiles (r = 0.63) reported in our results without site-specific reprogramming. This aligns with prior multi-vendor studies demonstrating reduced inter-scanner variability with Pulseq-based implementations [26]. Importantly, portability of the pulse sequence is only one component of standardized layer-fMRI: reconstruction, preprocessing, quality-control thresholds, and layerification steps can each influence laminar profiles. For this reason, we release reconstruction and analysis scripts alongside the Pulseq sequence file and emphasize the value of versioned software releases, standardized parameter templates, and transparent quality-control criteria for future multi-site deployments.

Pulseq also improves technical accessibility by lowering barriers to sequence development, by allowing sequences to be authored in MATLAB or Python and executed on supported systems without recompiling proprietary code. Graphical programming environments and client–server frameworks further facilitate interactive design and parameter exploration [28]. Consistent with this, our 5.0 T implementation benefited from rapid prototyping of reference and accelerated VASO acquisitions with matched parameters, reducing the need for low-level vendor-specific sequence programming when iterating sequence design. Together, published evidence and our experience support the view that Pulseq can accelerate development while promoting the standardization, portability, and accessibility essential for the widespread adoption of layer-fMRI.

### The M1 double-peak laminar signature reflects established cortical microcircuitry

The double-peaked VASO activation pattern observed at 5.0 T is consistent with established models of M1 laminar organization and supports the biological interpretability of our layer-resolved signals. M1 is an agranular cortex with pronounced separation of input and output pathways: superficial layers (Cytoarchitectonic layer II/III) primarily integrate converging inputs from premotor, somatosensory, and thalamocortical sources, whereas deep layers (specifically layer V) house large pyramidal neurons that generate the major corticospinal output [29–32]. This functional segregation is consistent with our VASO profiles. Quantitatively, the separation between superficial and deep peaks (∼5.60 depth bins) and the deep-to-superficial amplitude ratio (0.48 ± 0.18) summarize this laminar signature, with the superficial peak likely reflecting intracortical integrative processing and the deep peak corresponding to motor output generation. These quantitative descriptors can facilitate biological interpretation and cross-session comparisons under matched depth-sampling conventions. This interpretation is further supported by classical electrophysiological findings showing that corticospinal neurons in layer V of M1 increase firing during finger movements [33–35]. The detection of both peaks is enabled by VASO’s microvascular specificity [16], as evidenced by our observation that BOLD profiles showed only superficial dominance (7.47% ± 3.48%) whereas VASO resolved both superficial (4.81% ± 2.00%) and deep (2.10% ± 0.66%) components. Together, these findings position 5.0 T VASO as a robust tool for non-invasively characterizing the directional information flow within human cortical microcircuits.

### High efficiency and robust reproducibility of 5.0 T layer-fMRI facilitate the translation to clinical neuroscience

Building on the practical balance of sensitivity and accessibility described above, our results indicate that clinically feasible scan times and robust reproducibility metrics are achievable for laminar VASO in M1 at 5.0 T. We demonstrate that a single 13-minute acquisition is sufficient to resolve stable, layer-specific profiles (r = 0.78 vs. multi-run averages). This temporal efficiency is a crucial translational advantage: unlike prolonged research protocols, a 13-minute acquisition significantly improves feasibility for participants with limited scan tolerance, including pediatric and elderly cohorts and some clinical research populations. Complementing this efficiency is robust reliability: high test–retest reliability (r = 0.80) and strong between-subject generalizability (r = 0.86) support the utility of this framework for longitudinal tracking. Finally, these practical features are coupled with the mechanistic specificity of the VASO signal. Unlike BOLD, VASO mitigates large draining vein bias [1, 6–8], improving layer specificity and supporting the dissociation of superficial and deep components relevant to mesoscale input–output neural circuit hypotheses.

In the sensorimotor domain, this dissociation is clinically relevant. Superficial layers (Cytoarchitectonic layer II/III) receive thalamocortical and corticocortical inputs, whereas deeper layers (V/VI) give rise to corticothalamic and corticospinal output signals. In stroke and peripheral nerve injury, longitudinal 5.0 T imaging could track the reorganization of these input versus output pathways, providing a link between layer-specific plasticity and motor recovery [36–39]. Similarly, in motor disorders such as amyotrophic lateral sclerosis and focal hand dystonia, distinguishing relative changes in superficial versus deep engagement can help evaluate circuit-level dysfunction and treatment effects [40, 41]. This mesoscale input–output circuit hypothesis may also extend to sensory maladaptation: for example, in phantom-limb pain, animal models suggest that synaptic strengthening on Cytoarchitectonic layer V neurons in the somatosensory cortex correlates with pain intensity [42, 43]. Resolving such layer-specific plasticity in humans could validate these mechanistic targets.

More broadly, the framework’s high efficiency enables the testing of these circuit hypotheses beyond sensorimotor cortices. As 5.0 T systems become more widely available, this open and standardized framework could facilitate high-throughput investigations in association cortices and the hippocampus [44–48]—regions critical for neuropsychiatry and neurodegenerative conditions where large sample sizes are essential to overcome patient heterogeneity. In summary, by delivering a microvascular-specific readout of laminar brain activity that is standardized, reproducible, and feasible within a clinical timeframe, 5.0 T layer-fMRI establishes a viable ecosystem for translating mesoscale circuit models into robust clinical biomarkers.

### Technical considerations

Our protocol choices build on prior VASO literature while accounting for 5.0 T readout constraints. Prior work at 3.0 T demonstrated that, while BOLD weighting increases with TE, VASO contrast remained essentially invariant across 12–48 ms once BOLD correction was applied [14]. At 7.0 T, VASO protocols with TE in the 15–25 ms range and spatial resolutions between 0.8–1.5 mm have reported stable CBV sensitivity and effective mitigation of BOLD contamination [6, 8, 20, 49, 50]. Our choice of TE = 28.6 ms at 5.0 T accommodates hardware constraints while maintaining robust layer-specific CBV contrast and reducing residual BOLD contamination.

Beyond sequence optimization, ensuring the validity of the results requires rigorous analytical controls. To rule out the risk of circular analysis during ROI selection, we performed an additional independent validation analysis: the ROI was defined using the first VASO run of Visit 1, while laminar percent-signal-change profiles were extracted from the remaining runs. These independently derived profiles (Fig. S4) recapitulated the canonical double-peak pattern observed in the main results, confirming that our findings represent robust laminar signals rather than artifacts of ROI selection bias.

In addition to analytical rigor, precise depth conventions are essential for cross-study comparison. In our convention, depth-bin indices increase from the pial (CSF–GM) boundary toward the GM–WM boundary (bin 1 = most superficial; bin 11 = deepest). Across the 11 equidistant depth bins, both superficial and deep VASO peaks at 5.0 T appeared spatially shifted by approximately one depth bin towards the pial surface compared to canonical 7.0 T laminar patterns [1]. This spatial displacement is attributable to differences in cortical boundary definitions: our manually defined ROI included the pial surface (CSF-GM interface), whereas many 7.0 T protocols exclude the superficial 10-15% of cortical depth to reduce signal contributions from pial vessels. Crucially, while this results in a systematic superficial shift of absolute peak positions, the characteristic double-peak pattern and the relative separation of peaks (∼5.60 depth bins) remain preserved across field strengths, underscoring that laminar profile comparisons across studies must account for explicitly stated boundary conventions.

### Conclusions

This study establishes an open-source, Pulseq-based 3D-VASO layer-fMRI framework at 5.0 T, integrating a vendor-agnostic acquisition with an end-to-end reconstruction and analysis pipeline. At matched sub-millimeter resolution, benchmarking against a vendor-provided 2D-VASO protocol showed substantially expanded slab coverage (∼1.82×) and higher tSNR (∼1.50×). Using a finger-tapping paradigm in M1, this framework reliably resolved the canonical VASO double-peak laminar signature and demonstrated robust agreement across repeat visits, across participants, and across two independent imaging sites. Group-averaged profiles closely matched established 7.0 T laminar signatures, and even single 13-minute acquisitions preserved the canonical double-peak pattern, enabling higher-throughput study designs. By openly releasing this complete framework including the pulse sequence, reconstruction pipeline, and analysis scripts, we deliver a standardized, vendor-agnostic solution that enables reproducible and clinically feasible layer-fMRI, democratizing access to mesoscale human laminar mapping.

## Materials and Methods

### Participants

A total of 11 healthy young adults (aged 22–33 years; mean ± SD = 24.64 ± 3.17 years; 7 females) participated in this study. Ten participants were recruited from the college student population at the primary imaging site (Site 1), and one additional participant was recruited from the local community at an independent site (Site 2) for cross-site portability validation. All participants were right-handed, had normal or corrected-to-normal vision, and self-reported no history of neurological or psychiatric disorders. None were taking medications known to affect brain function. Written informed consent was obtained from each participant prior to study enrollment. This study was approved by the Research Ethics Review Committee of ShanghaiTech University. A schematic overview of this study is shown in Fig 1A.

### Task paradigm

Participants performed a visually guided block-design finger-tapping task targeting the M1 hand-knob region. Each run consisted of an initial 31.5 s pre-stimulus baseline, followed by 12 alternating task and baseline blocks (Fig 1B). Blocks lasted 31.5 s each, resulting in a total run duration of approximately 13 minutes. Baseline blocks consisted of a uniform black screen, and a 3 s onscreen countdown preceded each task block. During task blocks, participants viewed a video of a demonstrator repeatedly performing finger–thumb opposition and imitated the movements in real time by simultaneously tapping the index and middle fingers of the left hand against the thumb firmly at the demonstrated pace. Stimulus presentation was controlled using Experiment Builder (v2.6.11; SR Research Ltd., Ottawa, Canada) and synchronized to the start of each fMRI run via an MRI trigger pulse.

### Pulseq-based 3D-VASO sequence implementation and MRI data acquisition

#### MRI Hardware

Standard MRI safety screening was completed before each session to exclude contraindications. Brain MRI experiments were primarily conducted (n = 10) on a 5.0 T MRI scanner (uMR Jupiter, UIH, Shanghai, China) at Site 1. To assess the cross-site portability of the framework, one additional participant was scanned at Site 2 using an identical 5.0 T MRI scanner. Both scanners were equipped with a maximum gradient strength of 120 mT/m and a slew rate of 200 T/m/s. Imaging was performed using the standard vendor-provided 48/2-channel Rx/Tx head coil.

#### Pulseq 3D-EPI SS-SI-VASO implementation

A customized 3D echo-planar imaging (EPI) slice-saturation slab-inversion vascular space occupancy (SS-SI-VASO) sequence [6] was implemented using the Pulseq framework (v1.4.2; Figs 1C and 2A) [10] on the 5.0 T MRI scanner, with a vendor-provided Pulseq interpreter. The sequence design was adapted from a previously published Pulseq-based 3D EPI VASO sequence [25]. Sequence parameters were tested and optimized considering the scanner gradient coil performance and SAR constraints.

1. Inversion module. A time-resampled frequency-offset corrected inversion (TR-FOCI) pulse with 10.24 ms duration was positioned at the start of the sequence. The 100 ms duration between inversion pulse and the first excitation pulse was chosen to acquire the k-space center of the VASO imaging readout approximately at the magnetization nulling time (Mz-null) of once-inverted (non-steady-state) blood.
2. Excitation module. Sinc RF pulses with 3.7 ms duration were used for slice-selective excitation. Variable-flip-angle schemes were employed to control signal amplitude in the second phase-encoding direction of the 3D-EPI readout. This approach involved using an exponentially increasing flip angle for the first half of k-space, followed by a constant, higher flip angle for the second half.
3. Readout and acquisition parameters. The readout module comprised 20 kz-partitions, corresponding to 20 slices per slab. A fill time of 780 ms was inserted at the end of each pulse sequence repetition. Blood-nulled (VASO) and non-nulled (BOLD-weighted) images were acquired in an interleaved manner to enable BOLD-contamination correction of the nulled images. To support GRAPPA reconstruction for the accelerated acquisition, the VASO protocol paired (i) a fully sampled reference acquisition and (ii) an accelerated acquisition using an undersampled trajectory. Both acquisitions shared the same geometry and contrast timing, while the accelerated acquisition used in-plane parallel imaging and partial Fourier (see below). The following acquisition parameters were used for both the reference and accelerated sequences: field of view (FOV) = 192 × 192 × 24 mm³, voxel size = 0.8 × 0.8 × 1.2 mm³, effective repetition time (TR) = 4.5 s, echo time (TE) = 28.6 ms, first inversion time (TI₁) = 893.7 ms, and receiver bandwidth (BW) = 1080 Hz/pixel. For the accelerated sequence, additional parameters included: in-plane GRAPPA acceleration factor = 3 (effective echo spacing = 0.35 ms) and partial Fourier = 6/8.
4. Inversion time optimization for inflow suppression. TI₁ was defined as the duration between the start of pulse sequence and the acquisition of the k-space center of the blood-nulled readout. A critical constraint in this sequence is the inflow of non-inverted blood from outside the inversion slab, which introduces a cerebral blood flow (CBF)-weighted signal component that partially counteracts the intended CBV-weighted VASO contrast. Reducing TI₁ helps suppress this inflow contamination, as a shorter blood-nulling interval leaves less time for fresh blood to enter the imaging slab. One practical approach to shortening TI₁ is to allow for reduced inversion efficiency so that the magnetization null point occurs earlier [6]. Consequently, our choice of TI₁ = 893.7 ms was motivated by two considerations: (i) compensation for non-ideal inversion efficiency and partial T₁ recovery, and (ii) active suppression of non-inverted inflow contamination. This balance ensures that in our 5.0 T steady-state VASO implementation, robust CBV contrast is maintained while minimizing inflow-derived CBF bias.
5. Sequence validation. This 3D-VASO pulse sequence was implemented using the UIH ADEPT platform (Application Development Environment and Programming Tools; UIH, Shanghai, China) with a Pulseq interpreter. Sequence waveforms were visualized to confirm the intended pulse design. The gradient waveforms and k-space trajectory were exported and verified using KomaMRI (v0.9.0) [51]. To validate the end-to-end framework, a 3D brain phantom was used to solve the Bloch equations, generating raw k-space data in ISMRMRD format. These simulated data were then processed using the standard reconstruction pipeline (described in “*3D-VASO reconstruction and image quality benchmarking*”) to confirm image fidelity prior to scanner deployment.

#### MRI acquisition protocol

At Site 1, data collection followed a multi-visit design incorporating two independent VASO visits for test–retest reliability assessment. Each VASO visit consisted of two to four 3D-VASO fMRI runs and two structural MRI runs: one covering the whole brain and the other matched to the FOV of the 3D-VASO fMRI slab. At Site 2, the participant completed a single VASO visit using the same protocol as Site 1. All fMRI runs used the same task paradigm described above. The detailed protocol and acquisition parameters are as follows:

1. 3D-VASO fMRI runs for CBV-weighted layer-fMRI acquisition. To characterize layer-specific activity in M1, CBV layer-fMRI scans were acquired using the customized Pulseq-based 3D-EPI SS-SI-VASO sequence with the following parameters: FOV = 192 × 192 × 24 mm³, matrix size = 240 × 240 × 20, voxel size = 0.8 × 0.8 × 1.2 mm³, effective TR = 4.5 s, TE = 28.6 ms, TI₁ = 893.7 ms, BW = 1080 Hz/pixel, in-plane GRAPPA acceleration factor = 3, and partial Fourier = 6/8. A total of 175 pairs of blood-nulled and BOLD-weighted volumes were acquired for each run. The VASO imaging slab (Fig 1D) was positioned to center the M1 hand knob within the slice stack, guided by the structural images (described below). In eight participants, a BOLD-only fMRI run (described below) was also acquired to further guide the localization of the hand knob region. Each 3D-VASO fMRI run lasted 13 min 9 s.
2. MP2RAGE structural MRI runs for high-resolution anatomical imaging (vendor sequence: *gre_fsp_plus*). Two variants were acquired: a whole-brain acquisition, and a slab acquisition with the same coverage as the 3D-VASO fMRI runs. The whole-brain MP2RAGE parameters were: FOV = 180 × 190 × 145.6 mm³, matrix size = 256 × 270 × 208, voxel size = 0.7 × 0.7 × 0.7 mm³, TR = 7.9 ms, TE = 2.9 ms, TI₁ = 700 ms, TI₂ = 2500 ms, flip angle₁ = 7°, flip angle₂ = 5°, BW = 300 Hz/pixel, and parallel imaging acceleration factor = 6, number of averages = 1. Each whole-brain run lasted 4 min 5 s. For the slab MP2RAGE, parameters were optimized during the initial phase of the study to improve structural image quality. Consequently, for the majority of sessions (16 out of 20), the protocol used number of averages = 5, TE = 2.8 ms, and TI₁ = 700 ms. Other parameters were: FOV = 192 × 192 × 24 mm³, matrix size = 240 × 240 × 20, voxel size = 0.8 × 0.8 × 1.2 mm³, TR = 7.9 ms, TI₂ = 2500 ms, flip angle₁ = 7°, flip angle₂ = 5°, BW = 300 Hz/pixel, and parallel imaging acceleration factor = 6. Each slab run lasted 4 min 10 s. For the first four sessions, an initial version of the protocol was used with number of averages = 1, TE = 2.9 ms, and TI₁ = 755 ms, and each lasted 45 s. Both versions provided sufficient anatomical contrast for slab positioning and hand knob localization.
3. BOLD-only fMRI runs as functional localizers (vendor sequence: *epi_bold*). These runs were acquired using a 2D gradient-echo multiband EPI sequence to assist in localizing the hand knob region. The acquisition parameters were: FOV = 176 × 176 mm², matrix size = 220 × 220, number of slices = 48, voxel size = 0.8 × 0.8 × 0.8 mm³, multiband factor = 2, TR = 2000 ms, TE = 25.4 ms, flip angle = 90°, BW = 1200 Hz/pixel, and in-plane GRAPPA acceleration factor = 3. A total of 375 volumes were acquired for each run. The task paradigm comprised a 30 s pre-stimulus baseline followed by 12 alternating task and rest blocks (30 s each). This acquisition covered the primary motor and premotor cortices surrounding the hand knob region. Each run lasted 13 min 17 s.
4. 2D-VASO fMRI runs for benchmarking (vendor sequence: *epi_fid_r*). Two 2D-VASO fMRI runs were acquired in a single participant. The acquisition parameters were: FOV = 160 × 160 mm², matrix size = 200 × 200, number of slices = 11, slice thickness = 1.2 mm, voxel size = 0.8 × 0.8 × 1.2 mm³, TR = 4.5 s, TE = 28.6 ms, TI₁ = 1300 ms, TI₂ = 3700 ms, flip angle = 71°, BW = 1080 Hz/pixel, and in-plane GRAPPA acceleration factor = 3. A total of 175 pairs of blood-nulled and BOLD-weighted volumes were acquired per run with the same slab center matched to the 3D-VASO fMRI runs. Each run lasted 14 min 9 s.

#### Physiological data acquisition

Throughout all scanning visits, physiological signals were recorded synchronously with MR acquisition. Respiration and pulse were monitored using the scanner-integrated pneumatic belt and finger pulse oximeter, respectively. At Site 1, surface electromyography (EMG) was recorded to monitor task compliance using a BIOPAC MP160 system equipped with an isolated digital interface (STP100D) and an EMG amplifier (EMG100C-MRI) with AcqKnowledge software (v5.0.5; BIOPAC Systems, Inc., Goleta, CA, USA). EMG signals were sampled at 2 kHz with an amplifier gain of 2000, synchronized to scanner triggers, and online-processed using a 50 Hz comb band-stop filter in AcqKnowledge. Two recording electrodes were placed on the medial aspect of the left forearm, aligned longitudinally along the muscle belly, with a ground electrode positioned on the lateral aspect of the same arm (Fig S5).

### 3D-VASO reconstruction and image quality benchmarking

#### Image reconstruction

Offline reconstructions were performed using custom scripts in MATLAB (R2025a; MathWorks, Natick, MA, USA). Raw data of the accelerated scans in ISMRMRD format were filled into k-space according to the sequence sampling trajectories generated with Pulseq. Reconstruction included ramp-sampling interpolation to account for the trapezoid gradient readout. To reduce eddy-current-induced Nyquist ghost artifacts, the three navigator lines acquired prior to the k-space center were used to correct zeroth- and first-order phase errors. GRAPPA reconstruction was performed with a kernel size of 3 × 3, calibrated using the paired fully sampled reference acquisition acquired immediately before each accelerated run [52, 53].

#### Distortion assessment

To serve as a geometric reference, the MP2RAGE images (TI₁ and TI₂) were combined to generate a bias-field-corrected enhanced T1-weighted image (eT1w). Background noise was removed from the eT1w image using a regularized combination of the TI₁ and TI₂ images [54]. Automated segmentation of the eT1w image was subsequently conducted using FreeSurfer (v8.1) to generate cortical surface boundaries (CSF/GM and GM/WM) [55]. For distortion assessment, reconstructed 3D-VASO data and the denoised eT1w image were manually aligned using ITK-SNAP (v4.2.2) [56], and visual inspection was performed in terms of VASO geometric fidelity at the tissue boundaries (Fig 2B).

#### Quantitative benchmarking (3D vs. 2D VASO)

Image quality was quantitatively compared between the Pulseq-based 3D-VASO and the vendor-provided 2D-VASO sequence using two metrics:

1. Slab coverage: The slab coverage ratio was defined as the number of slices acquired in the 3D-VASO divided by the number of slices in the 2D-VASO (in-plane coverage and slice thickness matched).
2. tSNR ratio: tSNR was calculated voxel-wise as the mean temporal signal divided by the temporal standard deviation. To compare the sequences, a “tSNR ratio” (Mean tSNR 3D / Mean tSNR 2D) was computed in the anatomically defined hand-knob region for quantifying the relative tSNR gain. Statistical comparisons of voxel-wise tSNR distributions were performed using the Mann-Whitney U test in OriginPro (v2024; OriginLab Corporation, Northampton, MA, USA).

#### Functional signal verification

To verify the detection of functional activity, signal time courses were extracted from the 3D-VASO data. Following standard preprocessing (detailed below in “*Data preprocessing and quality control*”), ROIs were defined by intersecting functionally activated voxels (t > 4; sign-flipped to represent CBV increases) with the lateral aspect of the hand knob in Brodmann area 4a (detailed below in “*ROI definition and laminar profile extraction*”). Signal percent-change time courses for VASO and BOLD were extracted and visually compared against the task paradigm to confirm robust task-evoked responses.

### Data preprocessing and quality control

The following preprocessing procedures were performed prior to statistical modeling and laminar profiling, using AFNI (v24.1.03) [57], SPM (v12; https://www.fil.ion.ucl.ac.uk/spm/), LayNii (v2.3.0) [58], and custom MATLAB scripts.

#### Functional localizer preprocessing and analysis (BOLD-only fMRI)

Functional localizer BOLD-only fMRI data were analyzed to identify task-evoked activation areas to assist localization of the hand knob region. The preprocessing and analysis steps included: (1) rigid-body motion correction using AFNI’s 3dvolreg, with all functional volumes aligned to the first volume; (2) 2D spatial smoothing using a Gaussian kernel with an in-plane full-width-half-maximum (FWHM) of 1.2 mm; and (3) statistical analysis using SPM’s general linear model (GLM). The experimental design was modeled as a block design, and the six head motion parameters were included in the GLM as nuisance regressors. Statistical maps were visualized at uncorrected p < 0.01 and p < 0.05, with cluster extent thresholds of 10 and 20 voxels, respectively. The above procedures were integrated into a rapid processing toolbox executed immediately after each run. The processing time was approximately 7 min per run (Workstation configuration: Intel i7-13700, 64GB RAM), enabling experimenters to make immediate decisions about the coverage of the subsequent 3D-VASO fMRI runs based on these task activation maps and the structural MRI images of the hand knob.

#### VASO preprocessing (3D SS-SI-VASO)

The image reconstruction of the 3D SS-SI VASO acquisitions yielded separate VASO and BOLD time series. Both time series were processed similarly to the preprocessing procedures described in [59], with the BOLD data providing auxiliary inputs for VASO processing: (1) Thermal noise was reduced using NORDIC denoising [60] (magnitude-only option, isotropic patch size = 12 × 12 × 12, NORDIC patch overlap = 2, phase filter width = 3, g-factor patch overlap = 2), applied to both VASO and BOLD time series. (2) Brain extraction was performed on the BOLD data using AFNI’s 3dAutomask. (3) Head motion was corrected using SPM12’s realignment, aligning all runs within each visit to the mean image of the first run via six-parameter rigid-body transformation. This step was performed on the VASO and BOLD time series independently. (4) Temporal upsampling was applied with a factor of two on both VASO and BOLD data. (5) BOLD correction was performed using the LN_BOCO algorithm to remove BOLD contamination from the VASO data [58], yielding CBV-weighted VASO time series. The preprocessing steps listed above were performed independently for each individual functional run.

#### tSNR-based quality control

tSNR maps were computed on the NORDIC-denoised VASO time series. Runs were excluded if more than 30% of voxels within the hand-knob region fell below a tSNR threshold of 7.5. Following this exclusion criterion, on average 90% of voxels within the hand-knob region across the final dataset had tSNR above 7.5.

#### Motion-based quality control

Data quality was also assessed using framewise displacement (FD) derived from the six head motion parameters [61]. FD was computed as the sum of absolute translational and rotational head movements between consecutive fMRI frames, with rotations converted to equivalent displacements on a sphere of 50 mm radius approximating the size of the human brain. Motion censoring was implemented at two levels: (1) individual time points with FD > 0.4 mm (half of the in-plane voxel size) were excluded from analysis, and (2) entire task blocks were excluded if they contained ≥3 time points exceeding this threshold. Furthermore, entire runs were excluded from the final analysis if more than 50% of their task blocks failed to meet these criteria. Applying these criteria to the 11 participants resulted in a final dataset of 58 retained runs.

#### EMG processing and task compliance assessment

EMG data were collected at 2 kHz to verify task compliance. Offline processing was performed using a custom MATLAB script. First, the DC offset was removed by subtracting the mean from the raw EMG signal. A band-pass filter (2–45 Hz) was then applied to facilitate discrimination between resting and finger-tapping states. The filtered signal was converted to its absolute value and smoothed using a low-pass filter (cutoff frequency: 0.5 Hz), yielding a continuous linear envelope *x*(*t*).

Maximum voluntary contraction (MVC) reflects the maximum level of muscle activation that can be achieved under participants’ effort. MVC served as a reference for normalizing EMG signals across participants and recording sessions. For each run, MVC was defined as the maximum of the EMG linear envelope. A visit-level MVC was then obtained for each participant by taking the maximum MVC across all runs within the same visit. Time points at which *x*(*t*) exceeded 10% MVC were classified as active. For each task block, the compliance score S_i_ was defined as the proportion of active time points:

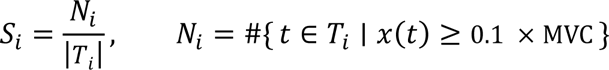

The compliance score for a run was computed as the average compliance score across all 12 blocks. The compliance score for a visit was computed as the average compliance score across all runs within that visit.

### ROI definition and laminar profile extraction

#### Statistical modeling and ROI definition

Statistical analysis was performed using a general linear model (GLM) via AFNI’s 3dDeconvolve. For each subject, preprocessed VASO data from all runs within each visit were analyzed jointly in a single GLM. The task design was modeled using a boxcar regressor convolved with a canonical hemodynamic response function (HRF). Six rigid-body motion parameters (three translations and three rotations) were included as nuisance regressors, and high-motion time points were censored using the run-specific censor vector. Baseline and low-frequency drifts were modeled separately for each run. Voxelwise t-maps were computed from this model and used for ROI definition. Because VASO signal intensity is inversely related to CBV, task activation results in negative regression coefficients. To facilitate visualization and ROI definition, the resulting t-statistics and beta values were sign-flipped (multiplied by −1) so that task-evoked CBV increases are represented as positive activation.

Prior CBV-weighted layer-fMRI studies of M1 commonly target the portion of the hand knob in Brodmann area 4a (BA4a) that expresses a characteristic superficial-and-deep (“double-peak”) laminar response during active movements [1, 21, 62]. Consistent with these established findings, we utilized this canonical signature as a functional landmark to identify the BA4a hand representation and to precisely distinguish the motor execution area (BA4a) from adjacent regions (e.g., somatosensory cortex) that often exhibit different laminar signatures. ROI definition followed a two-stage procedure combining anatomical constraints with functional selection (Fig 1E):

1. Anatomical hand-knob constraint. A cortical ribbon ROI was manually delineated on a single representative slice [14, 21] of the motion-corrected mean VASO image for the visit using ITK-SNAP, with the slab-matched structural image serving as an anatomical reference. The ribbon was restricted to the lateral hand-knob region within the precentral gyrus (approximating BA4a based on gyral and sulcal landmarks) [14]. The superior boundary of the ROI was placed at the CSF–GM interface and the inferior boundary at the GM–WM interface, positioned at the visually estimated midpoints of the respective transition zones (approximately 50% gray matter fraction). Lateral boundaries were drawn perpendicular to the local cortical surface to connect the superior and inferior borders [58].
2. Double-peak BA4a localization. Consistent with the known laminar distribution of corticocortical afferents and corticospinal efferents [29], to localize the BA4a hand-knob subregion where input and output laminae are anatomically segregated, we further refined the candidate ROI to a contiguous patch in which task-evoked VASO activation was present in both superficial and deep portions of the cortical ribbon (i.e., two depth-separated activation regions consistent with the expected double-peak BA4a signature). This double-peak criterion was used as an ROI localization rule for targeting BA4a, not as an inferential test of laminar response shape.

#### Laminar profile extraction

Geometric cortical layers were defined using LayNii’s LN2_LAYERS algorithm based on the equidistant principle [58]. The cortical depth was partitioned into eleven equidistant bins on the spatially upsampled grid (0.2 mm isotropic; Fig 1E). Laminar profiles were extracted via a multi-step averaging procedure: (1) For each voxel, VASO and BOLD signals were converted to percent signal change, using the mean of the 8 upsampled time points (equivalent to the 18 s immediately preceding each task block) as baseline; (2) Responses were averaged across task blocks and then across runs within each visit; (3) Layer-specific signals were computed by averaging values across voxels assigned to each depth bin. For visualization purposes, VASO profiles in Figs 3 to 6 were sign-inverted (multiplied by −1), as CBV increases produce signal decreases in VASO. However, all quantitative metrics were computed on the original (non-inverted) VASO percent signal change values using absolute response magnitudes as specified below.

Two metrics were computed to quantify laminar specificity: (1) Peak separation: This was defined as the absolute difference between the bin indices of the maximal VASO absolute percent signal change in superficial (bins 1–3) and deep layers (bins 7–9). (2) Deep-to-superficial ratio: This was defined as the maximum absolute response in deep layers (bins 7–9) divided by the maximum absolute response in superficial layers (bins 1–3). Both metrics were computed for each visit and averaged to obtain the group-level results for Site 1 (10 participants × 2 visits).

#### Independent validation analysis for controlling circularity

Because the double-peak signature was used as an ROI localizer, we performed an independent validation analysis to ensure that laminar profiles were not trivially imposed by ROI definition (i.e., to avoid using the same data for both ROI definition and signal extraction). For each participant, ROI boundaries and the corresponding depth-bin masks were generated solely from the first 3D-VASO run of Visit 1. These independently defined masks were then held fixed and applied unchanged to the remaining runs of Visit 1 to extract layer-specific VASO percent signal change using the same baseline definition and averaging procedure described above. This analysis ensured independence between the data used to define the ROI/depth masks and the data used for laminar profiling.

### Reliability, generalizability, and portability analyses

We quantified robustness of laminar VASO profiles at 5.0 T along three dimensions: cross-visit reliability, between-participant generalizability, and cross-site portability.

#### Cross-visit reliability (test-retest reliability)

To assess the reliability of laminar activation patterns within participants in Site 1, we employed complementary metrics to capture both across-depth laminar profile similarity and peak precision of specific layers.

1. Laminar profile similarity (Pearson’s r for cross-visit correlation): To quantify the consistency of the overall profile shape, the Pearson correlation coefficient (r) was calculated for each participant between the visit-level laminar profiles from Visit 1 and Visit 2, and then summarized across participants. This metric captures the general signal trend (rise and fall) across the cortical depths.
2. Peak-Position Reliability (ICC and κ_w_): While Pearson correlation captures the similarity of the laminar profile shape across depths, we additionally computed ICC and weighted kappa (κ_w_) to explicitly quantify the discrete spatial agreement of the peak locations. This ensures that the specific input and output layers are identified at consistent cortical depths across visits.

- Intraclass Correlation Coefficient (ICC (3,1)): A two-way mixed-effects model for single measures was used to assess the absolute agreement of peak bin indices [63, 64]. ICC values were interpreted as: < 0.2 (poor), 0.2–0.4 (fair), 0.4–0.6 (moderate), 0.6–0.8 (good), and > 0.8 (excellent) [65]. The formula is given by:

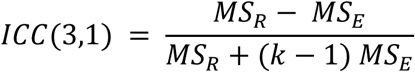 where *MS*_*R*_ is the mean square for rows (participants), *MS*_*E*_ is the mean square for error, and *k* is the number of visits (*k* = 2).
- Quadratic Weighted Cohen’s Kappa: To account for the ordinal nature of the depth bins (1–11), Cohen’s Kappa with quadratic weights was computed [66]. This approach assigns non-linear penalties to disagreements, placing greater weight on larger spatial deviations [67]. Interpretation followed standard guidelines: < 0 (poor), 0.00–0.20 (slight), 0.21–0.40 (fair), 0.41–0.60 (moderate), 0.61–0.80 (substantial), and 0.81–1.00 (perfect) [68]. The weighted kappa is defined as:

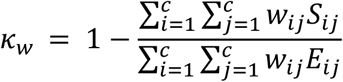 where *c* is the number of categories (11 depth bins), *S*_*ij*_ is the observed probability of Visit 1 classifying a peak in bin *i* and Visit 2 in bin *j*, and *E*_*ij*_ is the expected probability of agreement occurring by random chance (assuming statistical independence between visits). The quadratic weight *w*_*ij*_ is defined as:

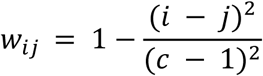

#### Between-participant generalizability

To assess robustness against cross-subject variability, the data in Site 1 (n = 10) were randomly divided into two independent subgroups (n = 5 per group). The consistency of the group-averaged laminar profiles was quantified using the Pearson correlation coefficients calculated between these two subgroups.

#### Cross-site portability

To evaluate portability, the identical sequence and task protocol were deployed at an independent validation site (Site 2, n = 1). Cross-site portability was quantified by correlating (Pearson correlation) the individual laminar profiles obtained from the participant in Site 2 with the group-averaged laminar profile obtained from Site 1.

### Statistical validation and benchmarking to published 7.0 T profiles

#### Statistical validation of the double-peak feature

To statistically validate the presence of the characteristic double-peak laminar pattern (i.e., whether signal in superficial and deep layers was significantly higher than in intermediate/boundary layers), linear mixed-effects (LME) models were implemented in MATLAB (MATLAB built-in function: fitlme). Laminar percent signal change values were extracted from the visit-level averaged laminar profiles (n = 20 visits at Site 1) and grouped into three categorical depth groups (*DepthGroup*): Superficial (bins 1–3), Deep (bins 7–9), and Reference (bins 4–6 and 10–11). The LME model was specified with percent signal change as the response variable, *DepthGroup* as a fixed effect, and nested random intercepts for subject and visit within subject. The model was formulated in Wilkinson notation as:

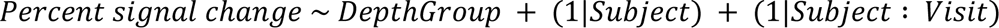

The “Reference” group (comprising the middle layers and GM-WM boundary) was treated as the reference level (intercept). Significance of fixed-effect coefficients was assessed for the Superficial and Deep groups relative to the Reference group, with p < 0.05 considered statistically significant.

#### Benchmarking to published 7.0 T profiles

To compare our group-averaged 5.0 T results (averaged across all 20 visits at Site 1) with previously reported laminar profiles acquired at 7.0 T [1], the group-averaged CBV-weighted (VASO) layer-specific fMRI response in M1 for the “tapping with touch” condition was digitized from the blue curve in Figure 3B of [1] using WebPlotDigitizer (https://automeris.io/). The original 7.0 T data were sampled at 20 equi-volume cortical depths [1]. The extracted profiles were subsequently resampled to 11 geometric layers using linear interpolation (MATLAB built-in function: interp1) to match the layering scheme of the current study for direct comparisons.

### Assessment of single-run robustness

To evaluate the sufficiency of a single 13-minute acquisition for resolving double-peak patterns and signal stability at the single-run level, a split-data validation analysis was performed. At Site 1, the first 3D-VASO run from each Visit 1 was treated as the “single-run” test dataset, while the average of all remaining runs within Visit 1 served as the high-tSNR reference (“ground truth”). Laminar profile similarity was calculated as the Pearson correlation coefficient (r) between the layer-specific signal profiles extracted from the first run and the high-tSNR reference profile. This calculation was performed at two levels: (1) Individual Level: Correlations were computed for each participant’s Visit 1 individually by comparing their single-run profile to their own reference profile. (2) Group Level: Data were first averaged across all participants to generate a “Group Mean Single-Run Profile” and a “Group Mean Reference Profile.” The Pearson correlation was then calculated between these two group-level laminar profiles.

Additionally, the spatial reliability of the two activation peaks in the single-run profiles was quantified relative to the reference profiles using ICC (3,1) and κ_w_, following the same methodology described above. This analysis determined whether the specific input/output layer peaks could be reliably localized using only 13 minutes of data.

## Data and Code Availability

The 3D-VASO Pulseq sequence file, image reconstruction and layer-fMRI analysis scripts, and example subject data are available at https://doi.org/10.12412/BSDC.1769409831.40001.

## Acknowledgments

This work was supported in part by funding from the Natural Science Foundation of Shanghai (24ZR1451500, Z.M.) and Shanghai Scientific Instruments and Chemical Reagents Project (24142201100, Z.M.). This work used the high-performance computing platform at ShanghaiTech University.

## Competing Interests

Haohao Peng, Yuhai Xie, and Jiayu Zhu are currently working as the employees of the United Imaging HealthCare Group Co., Ltd. and are receiving salary from this company. ShanghaiTech University has disclosed a provisional patent application CN202511776934.9 (filed by Z.M.).

## Author Contributions

**Conceptualization:** Zhiwei Ma.

**Data curation:** Yanting Zhu, Mengqi Jiang, Jianglian Chen, Fangyi Hao, Xin Li, Zhiwei Ma.

**Formal analysis:** Yanting Zhu, Mengqi Jiang, Jianglian Chen, Xin Li, Yiyun Qi, Zhiwei Ma.

**Funding acquisition:** Zhiwei Ma.

**Methodology:** Zhiwei Ma, Yanting Zhu, Mengqi Jiang, Jianglian Chen, Yiyun Qi, Yue Zhang, Haohao Peng, Yuhai Xie, Jiayu Zhu.

**Project administration:** Zhiwei Ma.

**Resources:** Zhiwei Ma.

**Supervision:** Zhiwei Ma, Jiayu Zhu.

**Writing–original draft:** Zhiwei Ma, Yanting Zhu, Mengqi Jiang, Jianglian Chen.

**Writing–review & editing:** Zhiwei Ma, Yanting Zhu, Mengqi Jiang, Jianglian Chen, Fangyi Hao, Xin Li, Yiyun Qi, Yue Zhang, Haohao Peng, Yuhai Xie, Jiayu Zhu.

