## Supplementary Information: Figure S1 to S5. for "Accessible and reproducible mesoscale fMRI at 5.0 T: A Pulseq-based open framework for human laminar mapping"

This file includes:

Figure S1 to S5: page 2 – 7

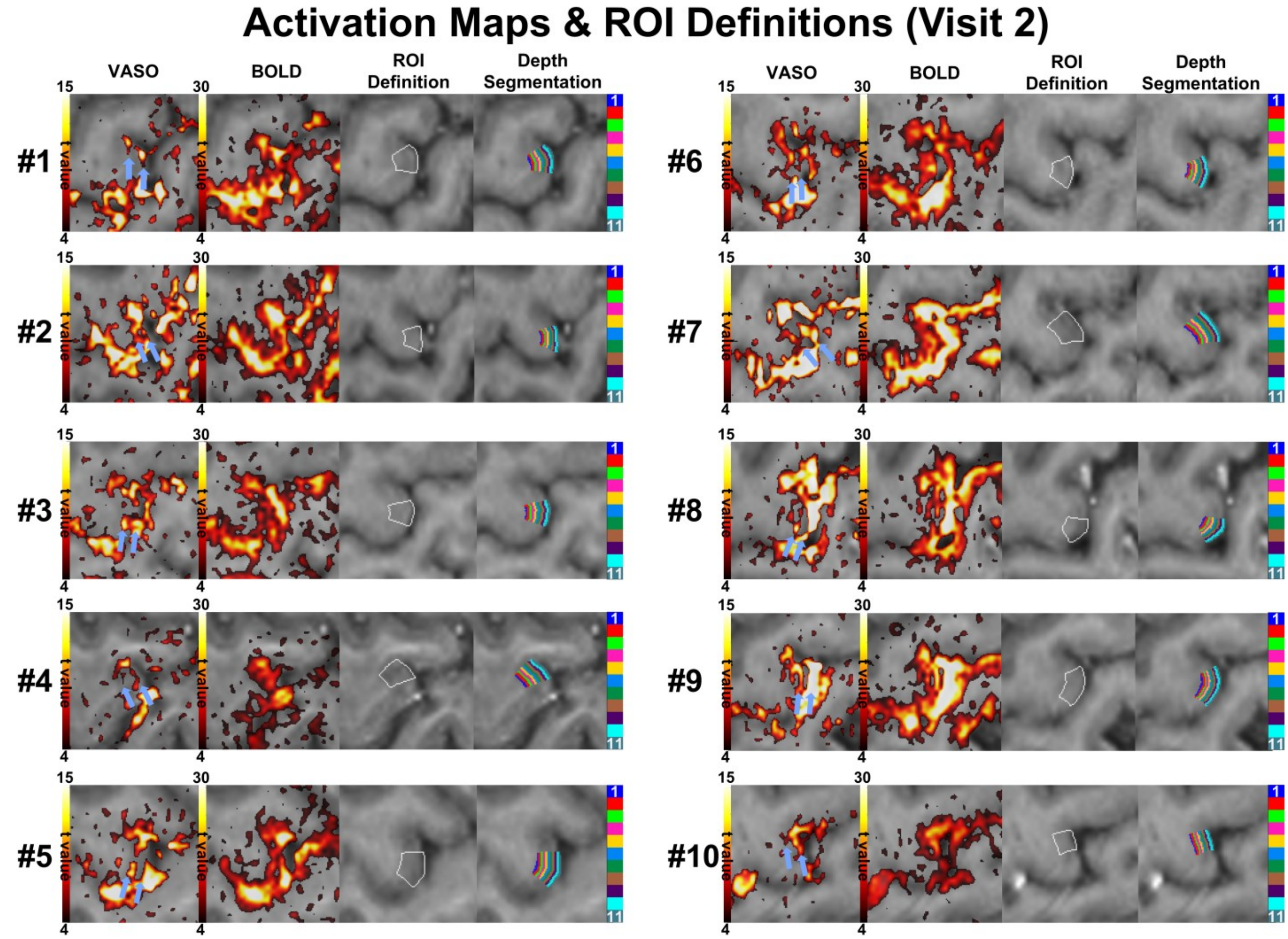


**S1 Fig. Individual-level activation maps and ROI definitions (Visit 2).** Columns display individual-level VASO (t > 4; sign-flipped) and BOLD (t > 4) activation maps, the ROI of the lateral hand-knob area (BA4a; white outline), and cortical depth segmentation (11 equidistant bins, L1–L11, from CSF to white matter). Blue arrows in VASO activation maps indicate the double-peak pattern (two depth-separated activation bands), whereas BOLD activation extends broadly into surrounding sulci.
**Abbreviations:** VASO, vascular space occupancy; BOLD, blood-oxygenation-level-dependent; ROI, region of interest; BA4a, Brodmann area 4a; CSF, cerebrospinal fluid.

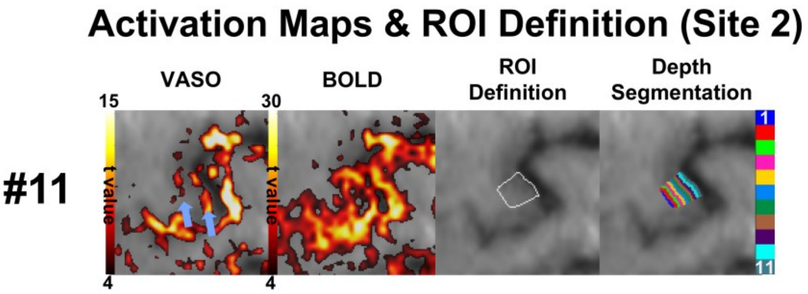


**S2 Fig. Individual-level activation maps and ROI definition (Site 2).** Columns display individual-level VASO (t > 4, sign-flipped; column 1) and BOLD (t > 4; column 2) activation maps, the ROI of the lateral hand-knob area (BA4a; white outline in column 3), and cortical depth segmentation (11 equidistant bins, L1–L11, from CSF to white matter; column 4). Blue arrows in VASO activation map indicate the double-peak pattern (two depth-separated activation bands), whereas BOLD activation extends broadly into surrounding sulci.

**Abbreviations:** VASO, vascular space occupancy; BOLD, blood-oxygenation-level-dependent; ROI, region of interest; BA4a, Brodmann area 4a; CSF, cerebrospinal fluid.

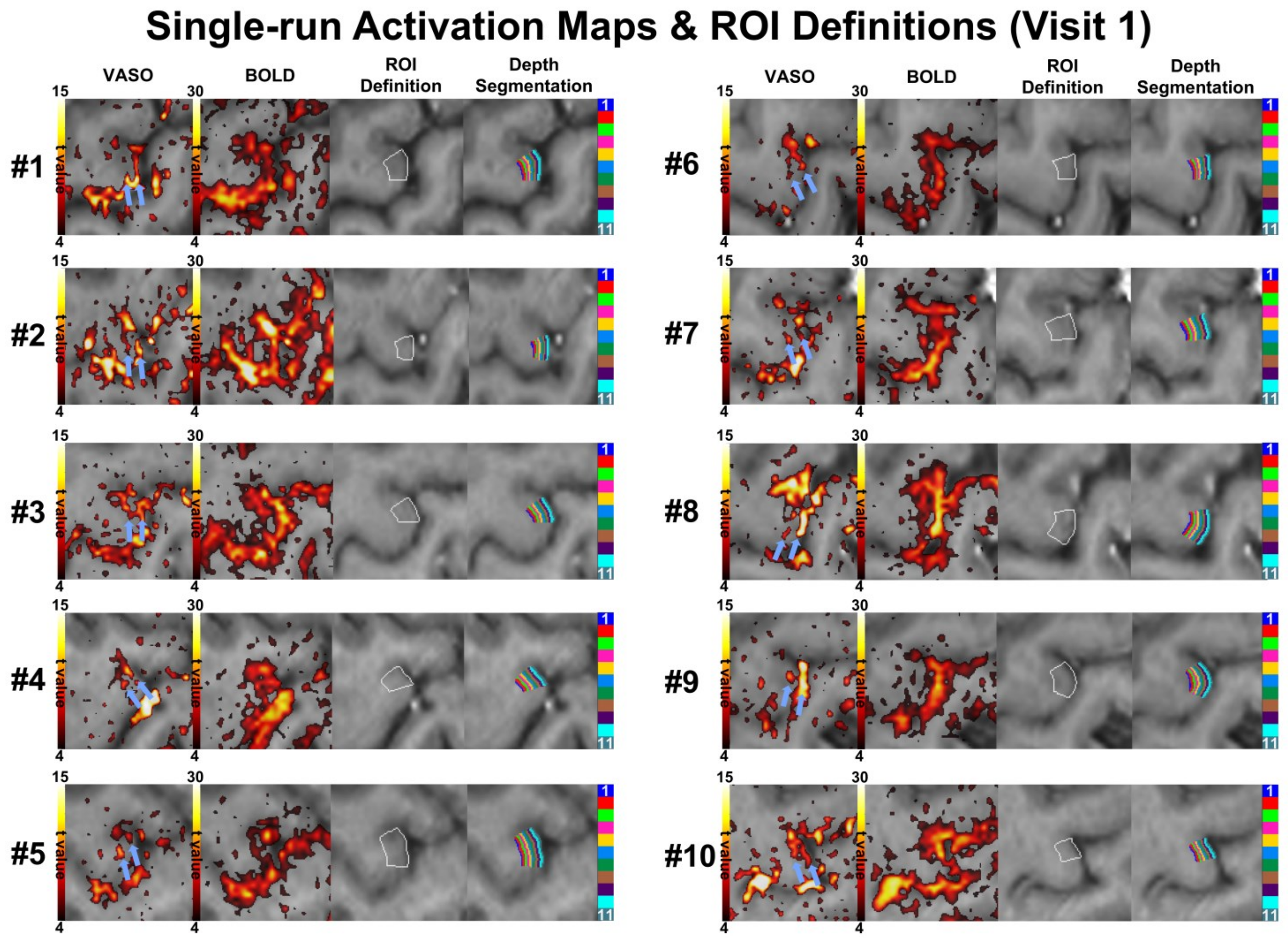


**S3 Fig. Individual-level single-run activation maps and ROI definitions (Visit 1).** Activation maps obtained solely from the first VASO run of Visit 1 for each participant. Columns display individual-level VASO (t > 4, sign-flipped) and BOLD (t > 4) activation maps, the ROI of the lateral hand-knob area (BA4a; white outline), and cortical depth segmentation (11 equidistant bins, L1–L11, from CSF to white matter). Blue arrows in VASO activation maps indicate the double-peak pattern (two depth-separated activation bands), whereas BOLD activation extends broadly into surrounding sulci.

**Abbreviations:** VASO, vascular space occupancy; BOLD, blood-oxygenation-level-dependent; ROI, region of interest; BA4a, Brodmann area 4a; CSF, cerebrospinal fluid.

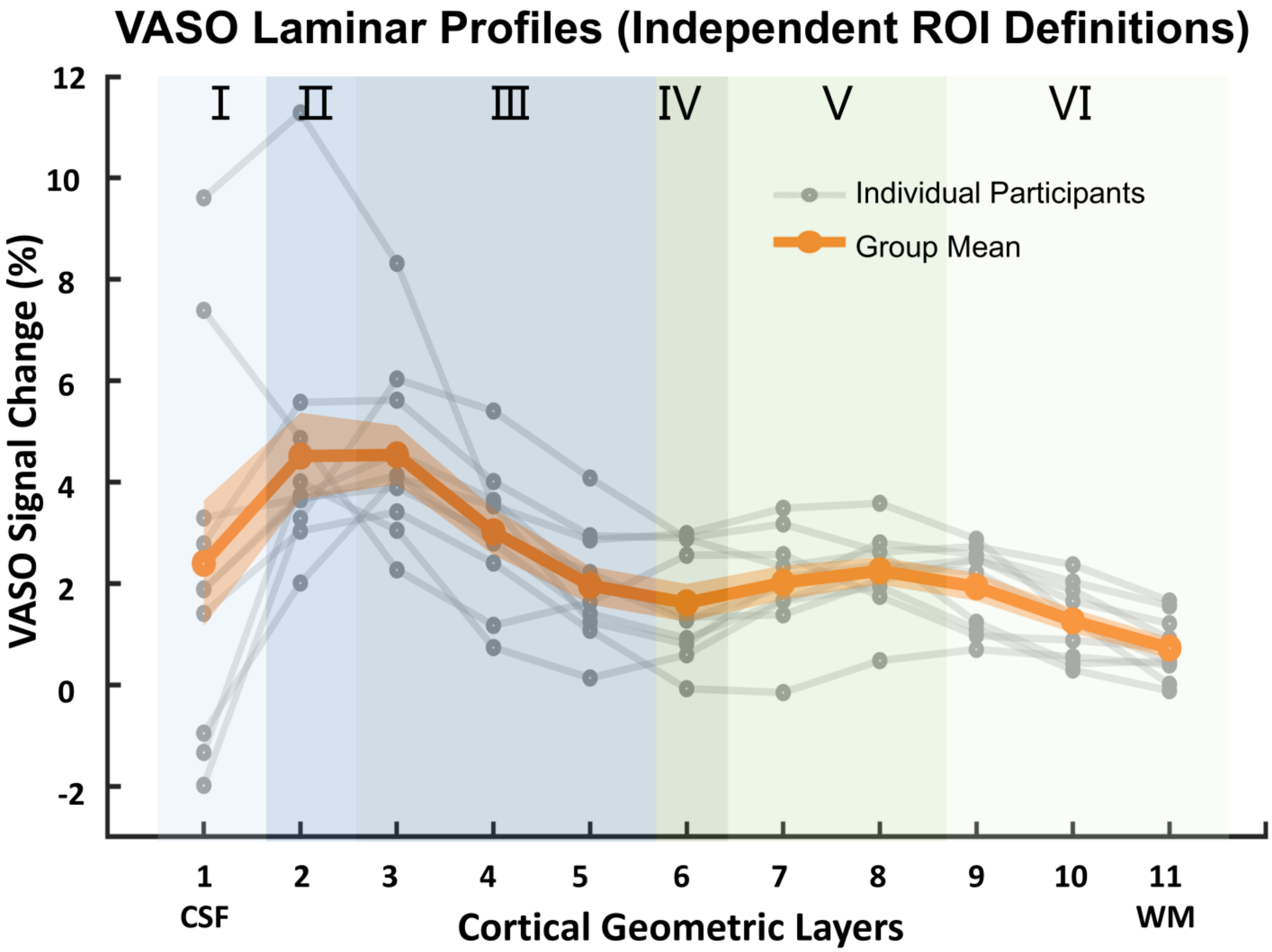


**S4 Fig. VASO laminar profiles computed with independent ROI definition.** ROI boundaries and cortical layering were generated solely from the first 3D-VASO run of Visit 1, while layer-specific percent signal changes were averaged from the remaining runs to control for circularity. Gray lines show individual participant profiles. The solid orange line represents the group-mean VASO response of Visit 1 based on this approach, and the shaded area indicates SE across participants. VASO profiles are sign-inverted for visualization. Background shading indicates cytoarchitectonic layers as in Fig 4.
**Abbreviations:** VASO, vascular space occupancy; ROI, region of interest; SE, standard error.

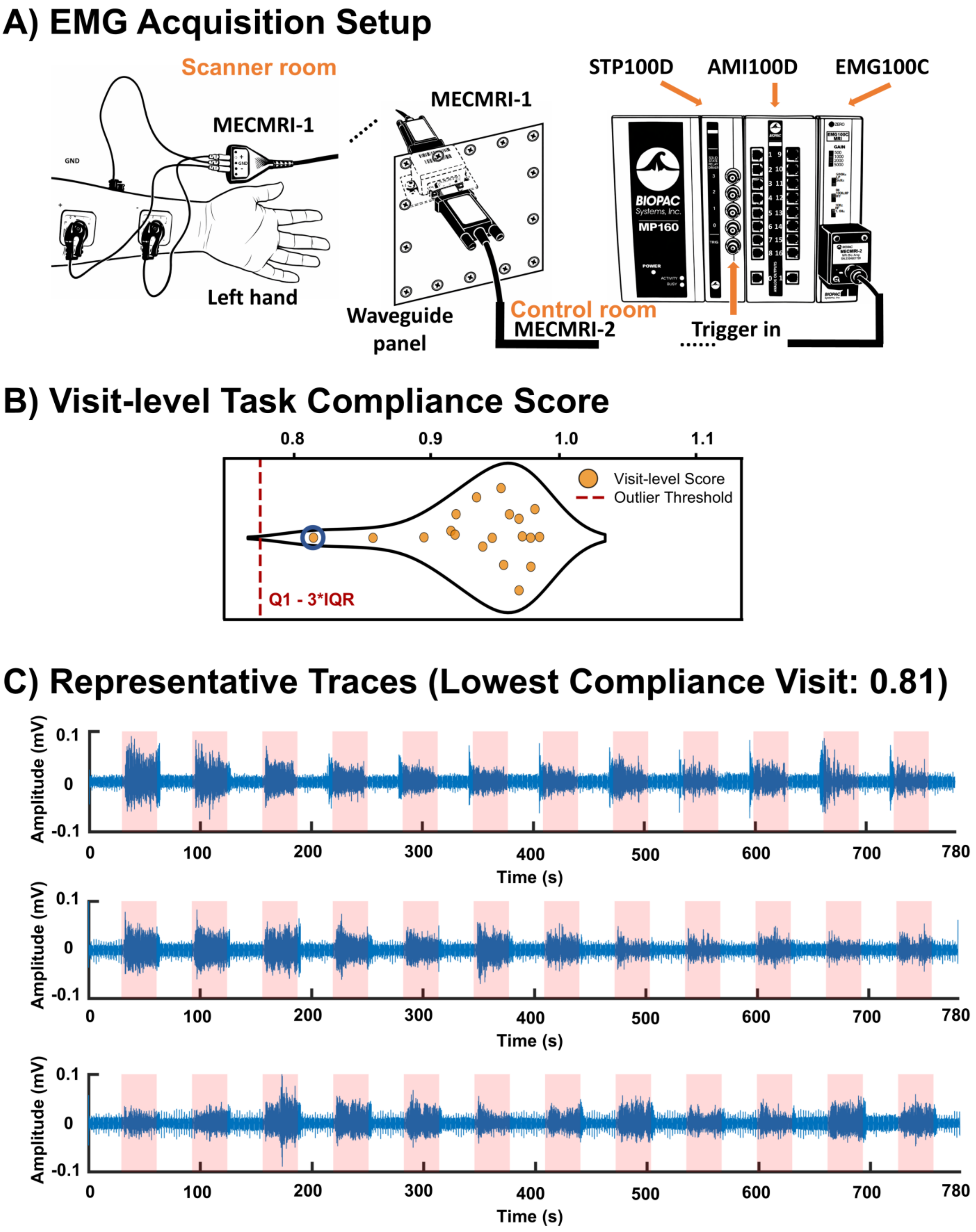


**S5 Fig. EMG acquisition setup and task compliance metrics. (A)** EMG acquisition setup. Surface EMG signals were recorded from the left forearm using two active electrodes and one ground electrode, routed through an RF waveguide, and transmitted to a BIOPAC MP160 system in the control room. Scanner triggers enabled synchronization with fMRI acquisition. **(B)** Visit-level task compliance scores

across participants. Each point represents the mean compliance score across all VASO runs within a visit (N = 11 participants; n = 19 data points total: Participant 7 Visit 1 EMG data were lost due to electrode detachment, and Participant 11 EMG data were not acquired). **(C)** Representative EMG traces from the visit with the lowest compliance score. The trace reveals clear task-modulated activity during each block despite having the lowest score, confirming robust compliance. The signal was preprocessed (DC-corrected, 2–45 Hz band-pass filtered) and divided by the amplifier gain (2000) to recover input voltages. Red shading indicates task blocks.
**Abbreviations:** EMG, electromyography; RF, radiofrequency; VASO, vascular space occupancy; DC, direct current.
